# Genome-wide CRISPR screens identify PTGES3 as a novel AR modulator

**DOI:** 10.1101/2025.05.29.656244

**Authors:** Haolong Li, James E. Melnyk, Becky Xu Hua Fu, Raunak Shrestha, Meng Zhang, Martin Sjöström, Siyu Feng, Jasmine A. Anderson, Wanting Han, Lisa N. Chesner, Hyun Jin Shin, Tatyanah Farsh, Humberto J. Suarez, Seema Nath, Jonathan Chou, Rajdeep Das, Emily A. Egusa, Jun Zhu, Aidan Winters, Ashutosh Maheshwari, Junjie T. Hua, Mohammed Alshalalfa, William S. Chen, Marsha Calvert, Elai Davicioni, Audrey Kishishita, Abhilash Barpanda, Tianyi Liu, Arun P. Wiita, Bradley A. Stohr, Javed Siddiqui, Bo Huang, Eric J. Small, Kevan M. Shokat, Peter Nelson, David A. Quigley, Elizabeth V. Wasmuth, Luke A. Gilbert, Felix Y. Feng

## Abstract

The androgen receptor (AR) is a critical driver of prostate cancer (PCa). To study regulators of AR protein levels and oncogenic activity, we created the first live cell quantitative endogenous AR fluorescent reporters. Leveraging this novel AR reporter, we performed genome-scale CRISPRi flow cytometry sorting screens to systematically identify genes that modulate AR protein levels. We identified and validated known AR protein regulators including HOXB13 and GATA2 and also unexpected top hits including PTGES3, a poorly characterized gene in PCa. PTGES3 repression resulted in loss of AR protein, cell cycle arrest, and cell death in AR-driven PCa models. PTGES3 is not a commonly essential gene, and our data nominate it as a prime PCa therapeutic target. Clinically, analysis of PCa data demonstrate that PTGES3 expression is associated with AR-directed therapy resistance. Mechanistically, we show PTGES3 binds directly to AR, forms a protein complex with AR in the nucleus, regulates AR protein stability *in vitro* and *in vivo* and modulates AR function in the nucleus at AR target genes. PTGES3 represents a novel therapeutic target for overcoming known mechanisms of resistance to existing AR-directed therapies in PCa.

## INTRODUCTION

The androgen receptor (AR) is a central driver of tumorigenesis and disease progression in prostate cancer^1,2^. The majority of prostate cancers express AR throughout the course of the disease^2–5^, and AR has been shown to promote transcriptional programs governing critical oncogenic phenotypes, such as proliferation, migration, and invasion^6^. Moreover, many key genes that drive prostate cancer (PCa), such as FOXA1 or HOXB13^7,8^, promote tumor progression by regulating how AR binds to or activates its target genes. Collectively, these findings highlight the importance of AR biology in this disease.

Multiple phase III clinical trials have demonstrated the benefit of AR-targeted therapies on patient survival^9–18^. AR signaling inhibitors (ARSI) are now the standard of care for locally advanced^12^, recurrent^13^, non-metastatic castration-resistant^9,17,18^, metastatic castration-sensitive^14–16^, and metastatic castration-resistant prostate cancer (mCRPC)^10,11^. However, aggressive PCa frequently escapes these therapies by reactivating AR signaling^19^. Recent analyses of clinical samples have demonstrated that the vast majority of mCRPC samples from patients resistant to ARSI agents abiraterone and enzalutamide exhibit robust AR/nuclear AR staining^20,21^. Therefore, identifying novel approaches to target AR is critical to improving survival outcomes for patients with AR-driven mCRPC.

We and other groups have identified tumor alterations that reactivate AR signaling during treatment with an ARSI^2,22–29^. These mechanisms of resistance include AR gene amplification^22,23^, AR enhancer amplification^2,24,25^, AR mutation^26^, AR genomic structural rearrangements^27^, AR splice variants^28^, and polymorphisms in androgen metabolism genes^29^. These alterations invariably result in increased expression or increased activity of the AR protein. Thus, strategies for reducing AR protein levels and AR activity represent a promising approach to overcoming known mechanisms of resistance in ARSI therapy-resistant AR-driven mCRPC. To identify such strategies, we sought to systematically and quantitatively identify genes that regulate AR protein levels with the goal of identifying next generation AR-targeting strategies for patients with aggressive AR-driven prostate cancer.

## RESULTS

### Establishment of an endogenous AR mNeonGreen2 (mNG2) fluorescent reporter in PCa cells

Accurate reporters of AR activity in prostate cancer cells are essential to developing the next generation of therapeutic approaches that target AR. Existing methods that fuse a full length fluorescent reporter protein to an endogenous or exogenously overexpressed target protein can affect target function, and are not ideal reporters^30,31^. We previously developed a split fluorescent protein tagging strategy enabling us to visualize and measure the expression of endogenous genes without antibodies^32^. This approach has advantages, including significantly reduced perturbation to the genomic locus and the protein-protein interactions of the target compared to traditional methods^32–34^. We set out to establish the first prostate cancer cell line models harboring an endogenous AR fluorescent reporter. To tag AR, we utilized a two-component split fluorescent protein tagging strategy that breaks the sequence of the monomeric NeonGreen (mNG2) protein between the tenth and the eleventh β-strand into two parts: mNG2_1-10 and mNG2_11, with mNG2_11 being a short, 16 amino acid peptide. Fluorescence arises only when the two protein fragments non-covalently bind, creating a fluorescent protein capable of serving as a reporter if fused to an endogenous gene. We knocked in mNG2_11 into the 5’ protein coding exon of *AR*, in the C42B cell model of aggressive prostate cancer, creating an N terminally tagged mNG2_11-AR fusion gene. We then stably expressed mNG2_1-10 in trans using a lentivirus and observed mNG2_1-10 complexes with mNG2_11-AR protein resulting in a bright fluorescent AR protein complex (**Fig.1a-d and Extended data fig.1**). We characterized multiple knock-in AR reporter clones (C42B^mNG2-AR^) by extensive genotypic and phenotypic approaches to confirm that tagged AR expression and stability as well as AR biological activity were unperturbed (**Fig.1c-i and Extended data fig.2**). Our approach allows us to rapidly and quantitatively measure endogenously expressed AR protein in live or fixed prostate cancer cells which should enable robust characterization of genetic or chemical perturbations that increase or decrease AR protein levels and localization.

**Figure 1.**
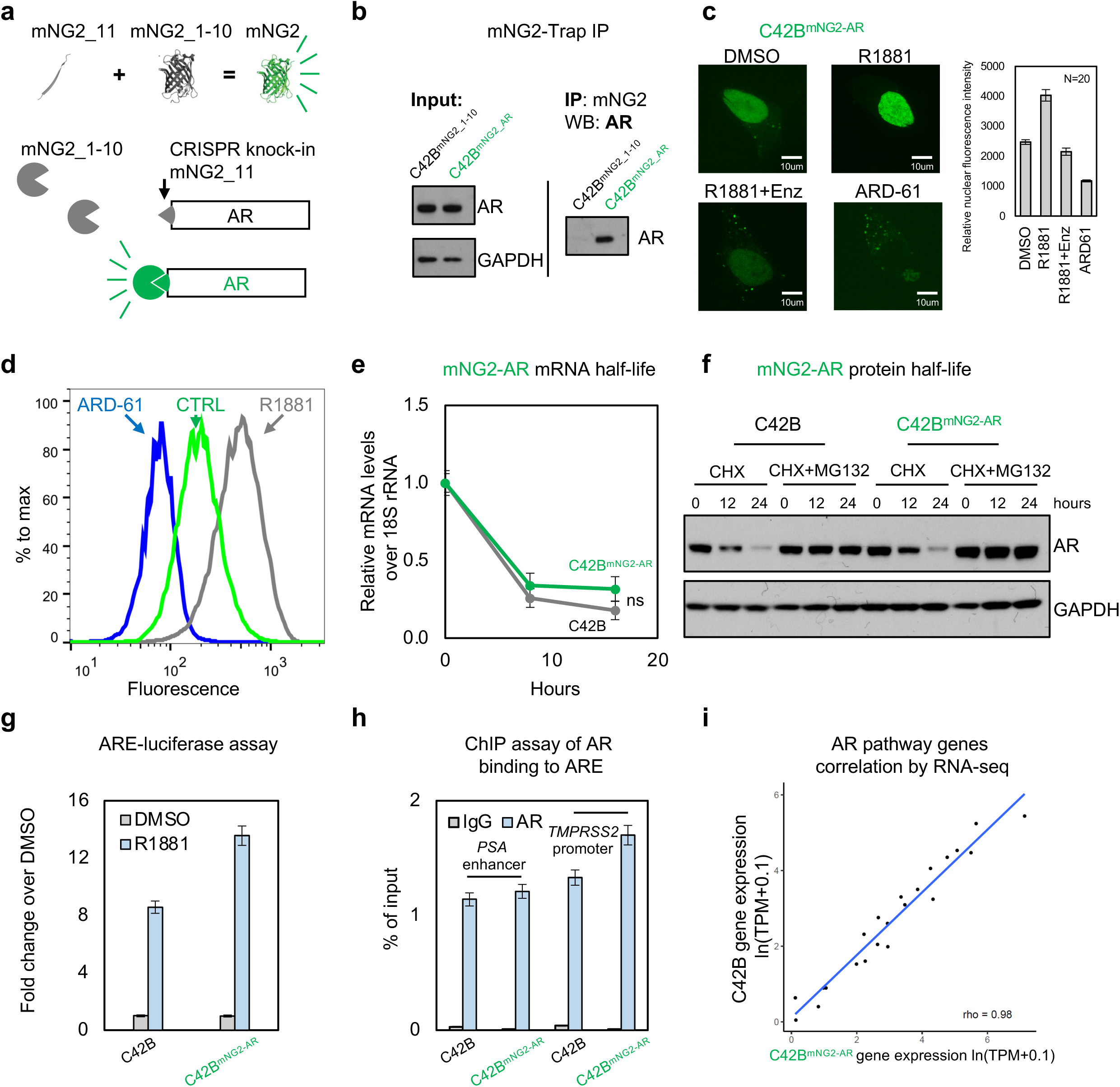
Establishment of an endogenous AR mNG2 fluorescent reporter in PCa cells. **a,** A diagram of the self-assembling mutated NeonGreen (mNG2) fluorescence protein system. The AR tagging design strategy: The 16 amino acid 11th β-strand of the fluorescent protein NeonGreen (mNG2_11) is knocked into the N-terminus of AR by CRISPR-Cas9. When the remainder of mNG2 (mNG2_1-10) is expressed in the same cell, it non-covalently binds to mNG2_11, producing a fluorescent protein. **b,** Immunoprecipitation (IP) experiments show the mNG2 antibody can pull down mNG2_1-10 and AR; protein levels were detected by western blotting. **c,** C42B^mNG2-AR^ cells were treated with DMSO, 10nM R1881 (androgen), 10nM R1881 plus 10μM Enzalutamide (Enz, anti-androgen), or 100nM ARD-61 (AR degrader, PROTAC) for 24h. Representative confocal microscopic images showed mNG2-AR localization (Green). Five images with total minimum 10 cells were collected per experimental condition. Nuclear fluorescence intensity was quantified using ImageJ. Error bars represent SEM. **d,** C42B^mNG2-AR^ cells were treated with DMSO (CTRL, green), 10nM R1881 (grey), or 100nM ARD-61 (blue). The mNG2-AR fluorescence from treated cells was analyzed by flow cytometry. Median Fluorescence Intensity (MFI) was calculated (n = 3; Mean ± SEM). **e,** C42B and C42B^mNG2-AR^ cells were treated with 1uM Actinmycin D for 0, 8 or 16 hours. RNA was collected to measure the AR mRNA levels using real-time PCR (n = 3 as biological replicates; Mean ± SEM; ns indicates no significant difference by two-way ANOVA analysis). **f,** C42B and C42B^mNG2-AR^ cells were treated with protein synthesis inhibitor Cycloheximide (CHX, 5μM) with or without proteasome inhibitor 5μM MG132 for 0, 12, 24 hours. AR protein levels were detected by western blotting. **g,** C42B and C42B^mNG2-AR^ cells were transfected with ARE firefly luciferase (ARE-Luc) then treated with DMSO (grey) or 10nM R1881 (blue). ARE-Luc over Renilla was normalized by control (n = 3 as biological replicates; Mean ± SEM). **h,** C42B and C42B^mNG2-AR^ cells were fixed by formaldehyde. ChIP experiments were performed using an IgG (grey) or AR antibody (blue). Precipitated DNA fragments were used as templates to amplify the PSA enhancer and TMPRSS2 promoter by real-time PCR (n = 3 as biological replicates; Mean ± SEM). **i**. Total RNA from C42B and C42B^mNG2-AR^ cells were collected for RNA-seq. Gene expression values were calculated as ln(TPM+0.1). Pearson correlation was calculated comparing AR pathway genes^84^ (n=22) between the two cell lines.

### Identification of regulators of AR using a genome-wide CRISPRi screen

We next set out to use our C42B^mNG2-AR^ cell lines to identify genes that regulate AR protein abundance. We first stably expressed a dCas9-KRAB fusion CRISPRi construct in C42B^mNG2-AR^ cells. Control experiments targeting TP53 and AR demonstrated robust CRISPRi gene silencing activity (**Extended data fig.3a-c**). We next performed genome-wide CRISPRi screens in two C42B^mNG2-AR^ clonal cell lines using fluorescence-activated cell sorting to identify genes whose repression would decrease or increase AR protein levels (**Fig.2a**). Genome-scale pooled genetic screens were performed by transducing C42B^mNG2-AR^ stably expressing dCas9-KRAB (C42Bi^mNG2-AR^) with a genome-wide CRISPRi library^35^ (**Extended data fig.3d**). Samples were collected, fixed with 3% PFA, and then sorted to isolate cells in the top and bottom quartile of AR fluorescent signal. We extracted genomic DNA and amplified the sgRNAs present and quantified sgRNA abundance in each sample by quantitative sequencing to identify genes that regulate AR levels. We confirmed that our screening strategy could robustly capture phenotypes induced by repression of genes such as AR or other commonly essential genes that induce strong growth phenotypes when repressed using time course fluorescent competition growth assays and by analysis of the abundance of commonly essential and non-essential sgRNA in our CRISPRi genome-scale screening data (**Extended data fig.3e-f**). We chose to solely focus on genes required for maintenance of AR levels, as such genes if targetable would be therapeutically relevant to prostate cancer therapy. As expected, sgRNAs targeting AR were strongly enriched in the low AR screen sample, meaning that repression of AR resulted in the greatest decrease in AR protein levels. We also identified other known AR regulators including *HOXB13*, *GRHL2*, and *GATA2* as top hits suggesting our screen robustly identifies genes required for AR protein abundance (**Fig.2b**). Top hits required for AR abundance include genes involved in chromatin transcription, RNA transportation, transcription factor, and post-translation regulation suggesting numerous biological processes influence AR protein levels. Top gene hits included protein complexes such as PAF and proteins such as HOXB13 and GATA2 known to interact directly or indirectly with AR^36^ (**Fig.2c**). Among them, FKBP4, PTGES3, UBE2I, HOXB13, and GATA2 showed evidence for physical subnetwork interactions with AR^37^. Gene Set Enrichment Analysis (GSEA) of the hit list revealed the androgen receptor signaling pathway as one of the top downregulated gene sets, suggesting AR protein levels may be regulated by AR target genes, as has been previously demonstrated for select examples such as FKBP4^38^ (**Extended data fig.4a**).

**Figure 2.**
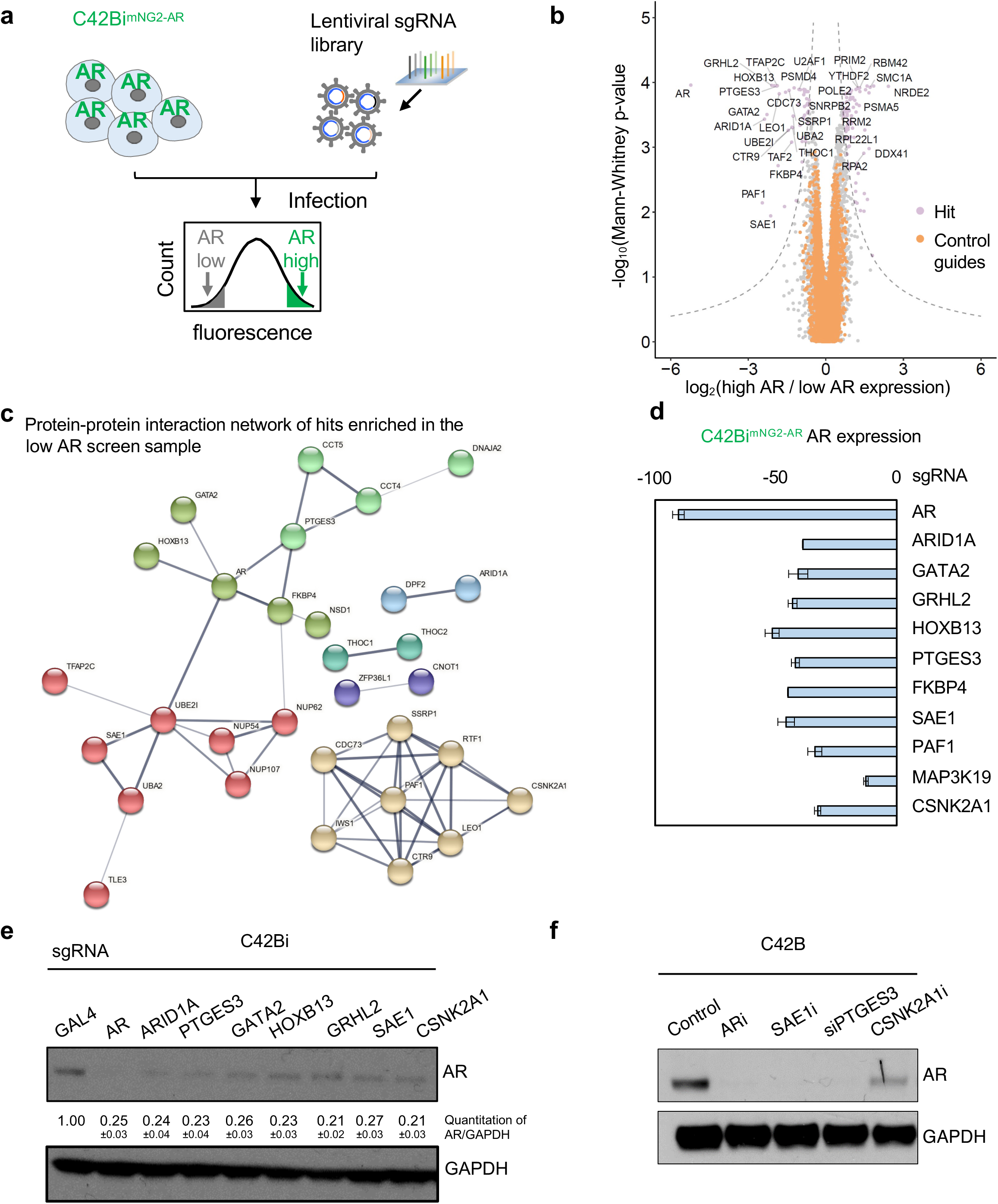
Identification of regulators of AR expression using a genome-wide CRISPRi screen. **a,** Schematic of the fluorescence-activated cell sorting (FACS) based genome-wide CRISPRi screening in C42Bi^mNG2-AR^ cells. **b,** A volcano plot of screening gene hits (light purple) and control guides (orange) were calculated by the established Weissman lab ScreenProcess protocol^51^. A comparable number of negative control (NC) genes were generated by randomly sampling 5 non-targeting sgRNAs (with replacement) and analyzed as true genes (n=∼19,000). Empirically derived thresholds (log_10_ p-value x phenotype z ≥ 13; dashed lines) were calculated as shown, using the NC gene distribution to derive the background standard deviation for z-score. Only a limited number of hits were labeled due to the size of the plot. **c,** Top positive AR regulator hits (n=51) protein interaction networks were analyzed by STRING. Genes without gene interactions among any of the top 51 hits were removed. Lines between the genes indicate a confident interaction. Line thickness indicates the strength of data support. Genes were also unsupervised clustered, indicated by their color. **d,** C42Bi^mNG2-AR^ were infected with sgRNA targeting control or indicated genes. After puromycin selection, Median Fluorescence Intensity (MFI) of mNG2-AR fluorescence from the treated cells was measured by flow cytometry. The percentage of AR expression was calculated as AR expression%=(MFI_C42B_-MFI_sg Individual gene_)/(MFI_C42B_-MFI_sgGAL4_)*100 (n = 3 as biological replicates; Mean ± SEM). **e,** C42Bi cells were infected with indicated individual sgRNAs. Cell lysates were collected after puromycin selection. AR and GAPDH levels were detected by western blotting. **f,** C42B cells were treated with 100nM ARD-61 (ARi), 5μM ML792 (SAEi), siRNA targeting PTGES3, or 10μM Slimitasertib (CSNK2A1i) for 24h. Cell lysates were collected. AR and GAPDH levels were detected by western blotting.

We validated individual top hit genes including *ARID1A*, *GATA2*, *GRHL2*, *HOXB13*, *PTGES3*, *FKBP4*, *SAE1*, *PAF1*, *MAP3K19*, and *CSNK2A1* in two C42Bi^mNG2-AR^ clonal cell lines demonstrating our screen results were reproducible and identified genes that modulate AR levels (**Fig.2d and Extended data fig.4b**). As a control we confirmed by quantitative RT-PCR (qRT-PCR) that CRISPRi knockdown of top hits decreased the mRNA expression of each targeted hit gene as expected (**Extended data fig.4c**). To extend our results beyond the C42Bi^mNG2-AR^ model, we confirmed repression of top hits identified in the screen decreased AR protein levels in parental C42B and LNCaP cells expressing dCas9-KRAB (hereafter C42Bi and LNCaPi), demonstrating the tagged mNG2-AR protein and unmodified AR protein are regulated by the same genes and further confirming the mNG2-AR model as a robust tool for studying AR biology (**Fig.2e and Extended data fig.4d**). We also validated our screen results using orthogonal genetic and chemical strategies, (ARD-61 for AR, ML-792 for SAE1, siPTGES3 for PTGES3, and CX-4945 for CSNK2A1). In each case, perturbation of candidate genes reduced AR protein levels (**Fig.2f**). Our results demonstrate genome-scale FACS based CRISPRi screens utilizing the C42B^mNG2-AR^ model robustly identified both known and unexpected genes that regulate AR levels.

### PTGES3 as a novel regulator of AR protein

We were intrigued by the identification of PTGES3 as a top regulator of AR levels. Using unbiased whole cell mass spectrometry-based proteomics analysis, we demonstrated significant downregulation of AR protein upon PTGES3 knockdown which confirms our western blotting results (**Extended data fig.5a**). Repression of PTGES3 did not alter *AR* mRNA levels as measured by qRT-PCR or RNA-seq, suggesting that PTGES3 regulates AR protein translation or protein stability (**Extended data fig.5c-d**). Knockdown of PTGES3 does not alter the mRNA or protein half-life of AR (**Extended data fig.5e-f**). As expected, our RNA-seq data showed that repression of PTGES3 disrupted AR activity resulting in decreased expression of AR target genes (**Fig.3a**). PTGES3 knockdown also decreased AR protein levels in other cell models of aggressive mCRPC including AR splice variant 7 expressing 22RV1 cells, AR gene amplified VCaP and AR-signaling inhibitor (enzalutamide) resistant MR49F cells (**Fig.3b**). These *in vitro* preclinical PCa models represent common clinically relevant AR statuses and our data leads us to speculate that a PTGES3 inhibitor could inhibit AR function in diverse high risk AR-driven mCRPC clinical scenarios.

**Figure 3.**
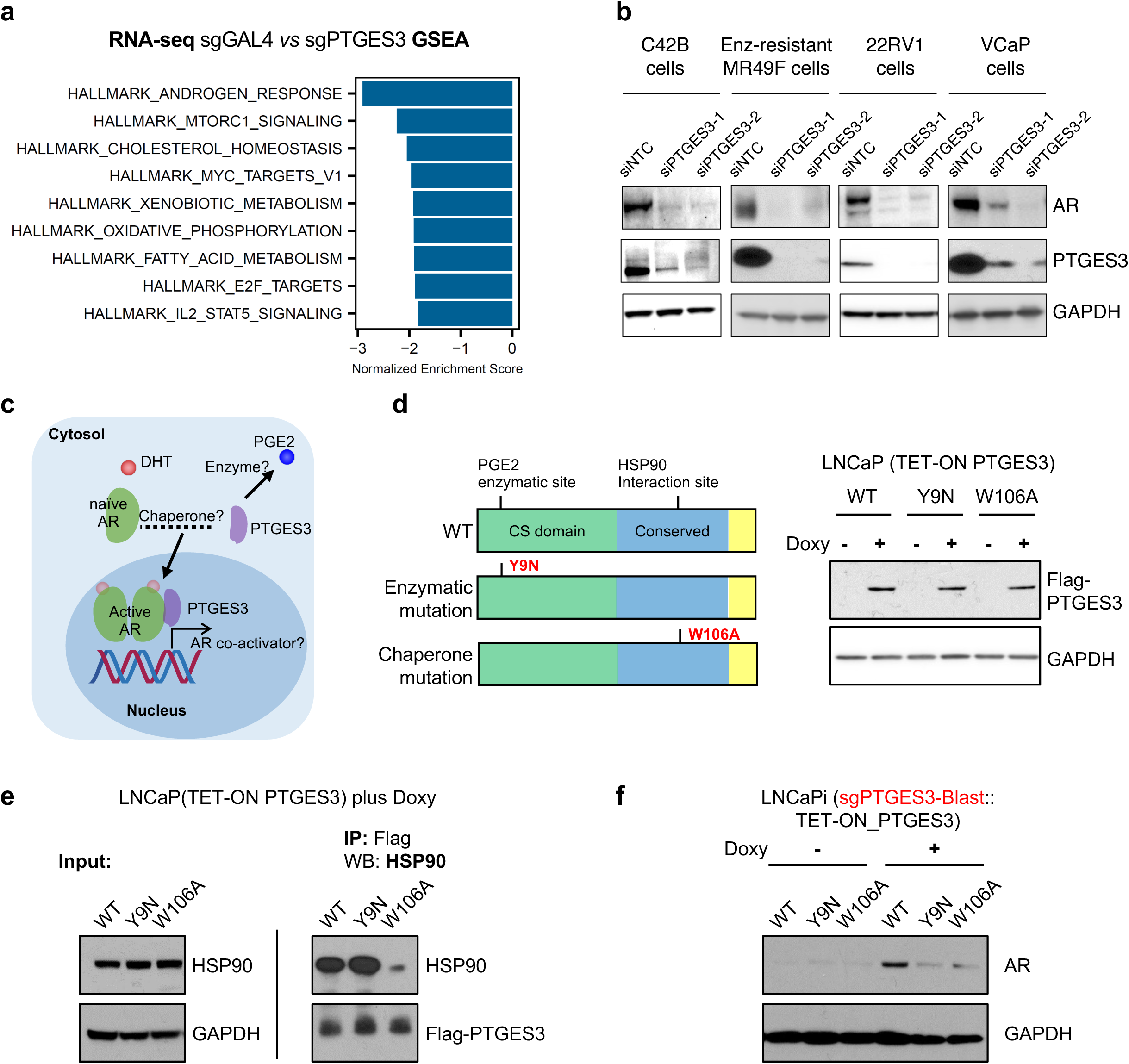
PTGES3 as a novel regulator of AR. **a**, LNCaPi cells were infected with sgRNAs targeting GAL4 (non-targeting control) or PTGES3. Total RNA was collected for RNA-seq (n = 2 as biological replicates). Gene Set Enrichment Analysis (GSEA) was performed. Hallmark_androgen_response is the top rank downregulated gene set. **b,** C42B, Enzalutamide resistant cell line MR49F, 22RV1, and VCaP cells were treated with siRNA targeting control or PTGES3. AR, PTGES3 and GAPDH levels were detected by western blotting. **c,** A diagram shows the potential dual functions of PTGES3 regulating AR. **d,** Left: Structures of the wildtype, enzymatic mutation (Y9N) and HSP90 mutation (W106A) isoforms of PTGES3. Right: LNCaP cells stably expressing a TET-ON inducible PTGES3 wildtype, Y9N mutation and W106A mutation were treated with or without 100ng/ml doxycycline. Flag-PTGES3 and GAPDH levels were detected by western blotting. **e,** LNCaP (TET-ON PTGES3) expressing inducible flag-tagged PTGES3 WT, Y9N mutation, or W106A mutation were treated with 100ng/ml doxycycline. Co-IP assays were performed using flag antibody. AR and HSP90 levels were detected by western blotting. **f,** LNCaPi cells were infected with sgPTGES3-Blast, then treated with DMSO or 100ng/ml doxycycline to overexpress the PTGES3 WT or Y9N or W106A mutant proteins. AR and GAPDH levels were detected by western blotting. Each western blot experiment was performed twice to determine reproducibility.

Having established that PTGES3 regulates AR levels in a variety of aggressive AR-dependent mCRPC cell models, we set out to decipher the mechanism by which PTGES3 supports AR protein levels and thus promotes AR signaling. In the absence of androgen ligands, AR protein is bound by protein chaperones such as HSP90 and localized in the cytosol^39^. Upon androgen stimulation, AR translocates into the nucleus, binds to androgen response elements and modulates gene expression of target genes^40^. As PTGES3 potentially regulates AR in a post-translational manner, we were interested in the protein function as well the localization of PTGES3. PTGES3 protein is reported to have multiple protein functions and to be localized to both the cytosol and nucleus^41,42^ (**Fig.3c**). PTGES3 is reported to have a cytosolic HSP90-dependent protein chaperone function for steroid nuclear receptors, but its activity in prostate cancer is poorly characterized^41,43,44^. PTGES3 is also reported to be a cytosolic prostaglandin synthase which converts Prostaglandin H2 (PGH2) to Prostaglandin E2 (PGE2)^45^; of note, other prostaglandin synthases in the same family as PTGES3 have been implicated in the production of androgens^46,47^. The proposed PGE2 enzymatic site is at the N terminus of the protein while the proposed HSP90 interaction motif is at the C terminus of the protein^41,45^.

To investigate the mechanism by which PTGES3 regulates AR, we confirmed that repression of PTGES3 or known co-chaperones of cytosolic AR, HSP90 or FKBP4, resulted in decreased AR protein levels as expected (**Extended data fig.6a**). Interestingly, repression of PTGES3 but not PTGES or PTGES2 reduced total AR protein levels (**Extended data fig.6b-c**). To dissect which of the two proposed PTGES3 protein functions regulates AR levels, we constructed lentiviral doxycycline inducible TET-ON C-terminal flag tagged constructs expressing wildtype PTGES3 (WT), PTGES3 with a PGE2 enzyme activity disrupting mutation (Y9N)^45^ and PTGES3 with a HSP90 interaction site mutation (W106A) (**Fig.3d**). We transduced LNCaPi (TET-ON PTGES3) with these constructs and created isogenic stable cell lines that overexpressed PTGES3 or PTGES3 mutant proteins upon addition of doxycycline (Doxy) (**Fig.3d**). We performed co-immunoprecipitation (IP) western blotting experiments on each PTGES3 construct and confirmed that the PTGES3 W106A mutation but not the Y9N mutation abrogates the interaction between PTGES3 and HSP90, indicating the Y9 site might not interact with HSP90 and suggesting the catalytic Y9 site could be critical for HSP90 independent function of PTGES3 **(Fig.3e)**. To test which of these PTGES3 mutations modulates endogenous AR levels, we knocked down endogenous PTGES3 in LNCaPi and then inducibly overexpressed the PTGES3 WT or W106A or Y9N mutant proteins. As expected, we observed that expression of wildtype PTGES3 rescued AR protein levels. Expression of either the Y9N or W106A mutant PTGES3 protein failed to rescue AR protein levels, suggesting that both sites/functions are required to maintain AR (**Fig.3f and Extended data fig.6d**). Additionally, we observed that loss of AR following PTGES3 knockdown is not rescued by inhibition of either the ubiquitin-proteasome or the lysosomal degradation pathway and is independent of the presence of androgen or enzalutamide (**Extended data fig.6e-g**).

### Nuclear PTGES3 facilitates AR mediated transcription

PTGES3 has also been reported to localize to the nucleus and a previous publication showed nuclear PTGES3 is a glucocorticoid receptor (GR) co-factor that modulates GR activity in the nucleus at GR response elements^48^. We set out to evaluate whether PTGES3 is localized to the nucleus in prostate cancer tumors. We measured PTGES3 localization by immunohistochemistry (IHC) using a validated PTGES3 antibody on a tissue microarray containing 120 prostate cancer tumor biopsies. PTGES3 was present at both cytosolic and nuclear locations with increased PTGES3 protein expression in more advanced/aggressive prostate cancer tumors (**Extended fig.7a**). Integrated analysis of PTGES3 localization with clinical outcomes data demonstrated patients with a high nuclear PTGES3 score had worse PSA-recurrent free survival, demonstrating that nuclear PTGES3 protein levels are correlated with a more aggressive prostate cancer phenotype (**Fig.4a**, HR=2.655, P log rank=0.0014).

**Figure 4.**
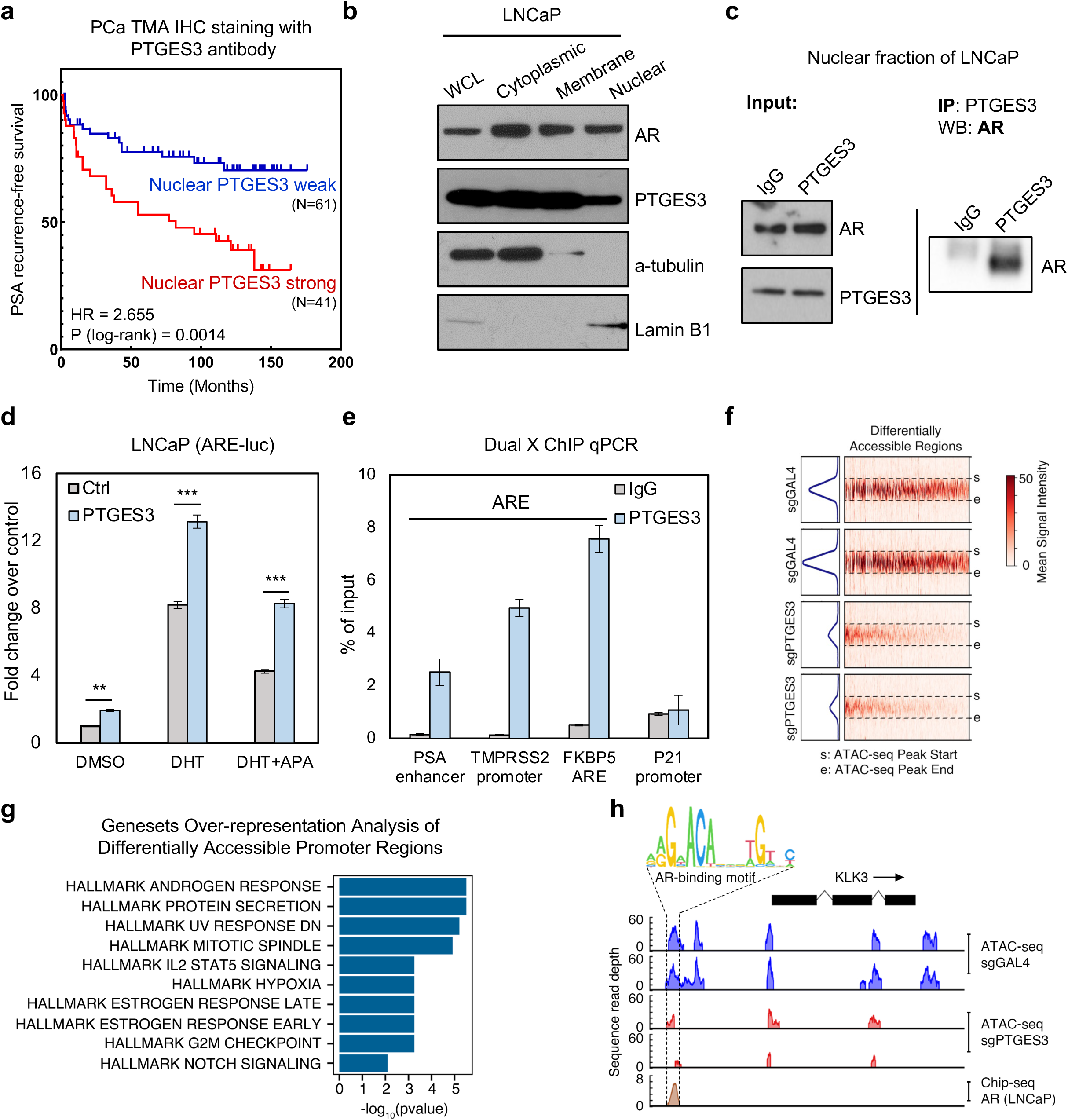
Nuclear PTGES3 facilitates AR mediated transcription. **a,** Immunohistochemistry was performed on human prostate cancer tissue microarrays (n=120). Cores were scored for intensity of nuclear and cytoplasmic PTGES3 staining. Kaplan Meier graph depicting relation between PSA recurrence free survival and nuclear PTGES3 staining in prostate cancer patients. Weak=score 0 or 1, Strong=score 2 or 3. **b,** Cell fractions from LNCaP cells were immunoblotted with indicated antibodies. **c,** Nuclear IP experiments were performed using IgG or PTGES3 antibody. AR protein levels were detected by western blotting. Each western blot experiment was performed twice to determine reproducibility. **d,** LNCaP cells containing ARE-luciferase reporter were transfected with pCDH-control vector (grey) or pCDH-PTGES3 (blue). Cells were treated with DMSO, 1nM DHT, or 1nM DHT plus 5μM Apalutamide (APA). Luciferase activities over Renilla were normalized with control (n = 3 as biological replicates; Mean ± SEM). Unpaired two-tailed t-test was used to determine statistical significance (**P < 0.01; ***P < 0.001). **e,** LNCaP cells were fixed sequentially by EGS and formaldehyde. Dual cross-linking ChIP experiments were performed using indicated antibodies. Precipitated DNA was used as a template to amplify the indicated genomic regions by real-time PCR (n = 3 as biological replicates; Mean ± SEM). **f,** LNCaPi cells were infected with sgGAL4 or sgPTGES3. Cells were collected for ATAC-Sequencing (n = 2 as biological replicates). The heatmap shows the differentially accessible ATAC-seq peak regions in sgPTGES3 as compared to that in sgGAL4. Each ATAC-seq peak region was scaled to the same size from the start to the end region. Flanking ±500bp regions are also shown. The top panel shows the corresponding ATAC-seq signal intensity. **g,** Genesets over-representation analysis of the differentially accessible ATAC-seq peaks (in sgPTGES3 as compared to that in sgGAL4) mapped to the promoter regions. The nearest genes to the promoter regions were selected for the over-representation analysis against the MSigDB Hallmark gene signatures. **h,** The figure shows ATAC-seq peak coverage in KLK3 gene body and flanking ±5kb region. sgGAL4 ATAC-seq peaks are colored in blue and sgPTGES3 ATAC-seq peaks in red. The upstream promoter region of KLK3 contains overlapping ATAC-seq peaks and AR Chip-seq peaks^48^. AR binding motif was identified in this region.

We further measured PTGES3 localization in the cytoplasm, membrane, and nuclear compartment of LNCaP cells. Western blot analysis demonstrated that PTGES3 was localized to both the cytoplasm and nucleus of LNCaP cells (**Fig.4b**). As expected, AR was localized in both the cytoplasm and nucleus while HSP90 was localized to the cytoplasm (**Fig.4b and Extended data fig.7b)**. To investigate whether PTGES3 modulates AR in the nucleus, we first tested whether PTGES3 interacts with AR in the nucleus by co-IP western. Our results demonstrated that PTGES3 physically interacts with AR in the nucleus (**Fig.4c**). Overexpression of PTGES3 significantly increased AR activity as measured by an Androgen Response Element (ARE) luciferase transcriptional response reporter activity assay, further supporting the hypothesis that PTGES3 expression promotes AR activity (**Fig.4d**). Consistent with a previous report^48^, among the transcriptional regulators p300, KAT2A and FOXA1, nuclear PTGES3 interacts with histone acetyltransferase KAT2A by co-IP western (**Extended data fig.7c-d)**. Knocking down KAT2A inhibited the enhanced AR activity observed upon overexpression of PTGES3 (**Extended data fig.7e)**. These evidence demonstrate PTGES3 potentially facilitates interactions with AR and transcriptional machinery in the nucleus.

PTGES3 is not predicted to bind directly to DNA. To test the hypothesis that a PTGES3/AR protein complex is localized to canonical endogenous AREs and AR target genes, we assessed PTGES3 genomic localization by indirect protein/DNA interactions by dual cross-linked IP followed by real-time PCR for DNA sequences of interest (Dual X ChIP). Our results demonstrated that PTGES3 is specifically enriched at AREs that are hallmark AR targets but not in non-AR binding region or control region (**Fig.4e**). Importantly, we observed that HSP90 is not enriched at hallmark AREs, further supporting the hypothesis that nuclear PTGES3 plays an HSP90-independent role in promoting AR activity **(Extended data fig.8)**. To test whether PTGES3 activity promotes chromatin accessibility and therefore potentially AR binding to regulatory elements, we performed Assay for Transposase-Accessible Chromatin using Sequencing (ATAC-Seq) comparing PTGES3 knockdown to controls. We observed repression of PTGES3 results in far more loss of open chromatin regions (OCRs) than gain of OCRs (**Fig.3f and Extended data fig.9a**). We then mapped the differentially accessible DNA regions to the nearest gene promoter and observed a strong enrichment for genes that are hallmark androgen response genes (**Fig.4g**). Integration of our ATAC-seq data with AR ChIP-seq data^49^ demonstrated that more than 80% of the differentially accessible ATAC seq peaks overlapped with AR ChIP-seq peaks (**Extended data fig.9b**). For example, knockdown of PTGES3 significantly reduced the chromatin accessibility at the canonical ARE of the *KLK3* promoter (**Fig.4h**). These data support the hypothesis that nuclear PTGES3 forms a protein complex with AR which is required for AR protein stability and transcriptional activity in AR-driven PCa and that this biology is distinguished from cytoplasmic AR co-chaperones such as HSP90, which in turn supports the notion that an anti-cancer strategy targeting PTGES3 will not phenocopy targeting HSP90.

### PTGES3 is a potential therapeutic target for AR-directed therapy resistant PCa

We next asked whether *PTGES3* expression is associated with PCa disease progression in mCRPC patient cohorts. We utilized a clinical-grade Affymetrix Human-Exon microarray to determine *PTGES3* expression levels in 641 prostatectomy samples from patients with high-risk prostate cancer. When we matched patients by clinicopathologic variables, including Gleason score, PSA, tumor stage, and nodal status, we determined that *PTGES3* overexpression was associated with significantly worse metastasis-free survival in patients who received adjuvant first-line androgen deprivation therapy (ADT) (with leuprolide or an GnRH agonist) after prostatectomy, but not in patients who did not receive adjuvant ADT after prostatectomy (**Fig.5a-b**). Furthermore, higher tumor expression of *PTGES3* was correlated with worse overall survival in mCRPC patients treated with a first-line AR targeted therapy (abiraterone acetate, enzalutamide, or apalutamide) (**Extended data fig.10a**).

**Figure 5.**
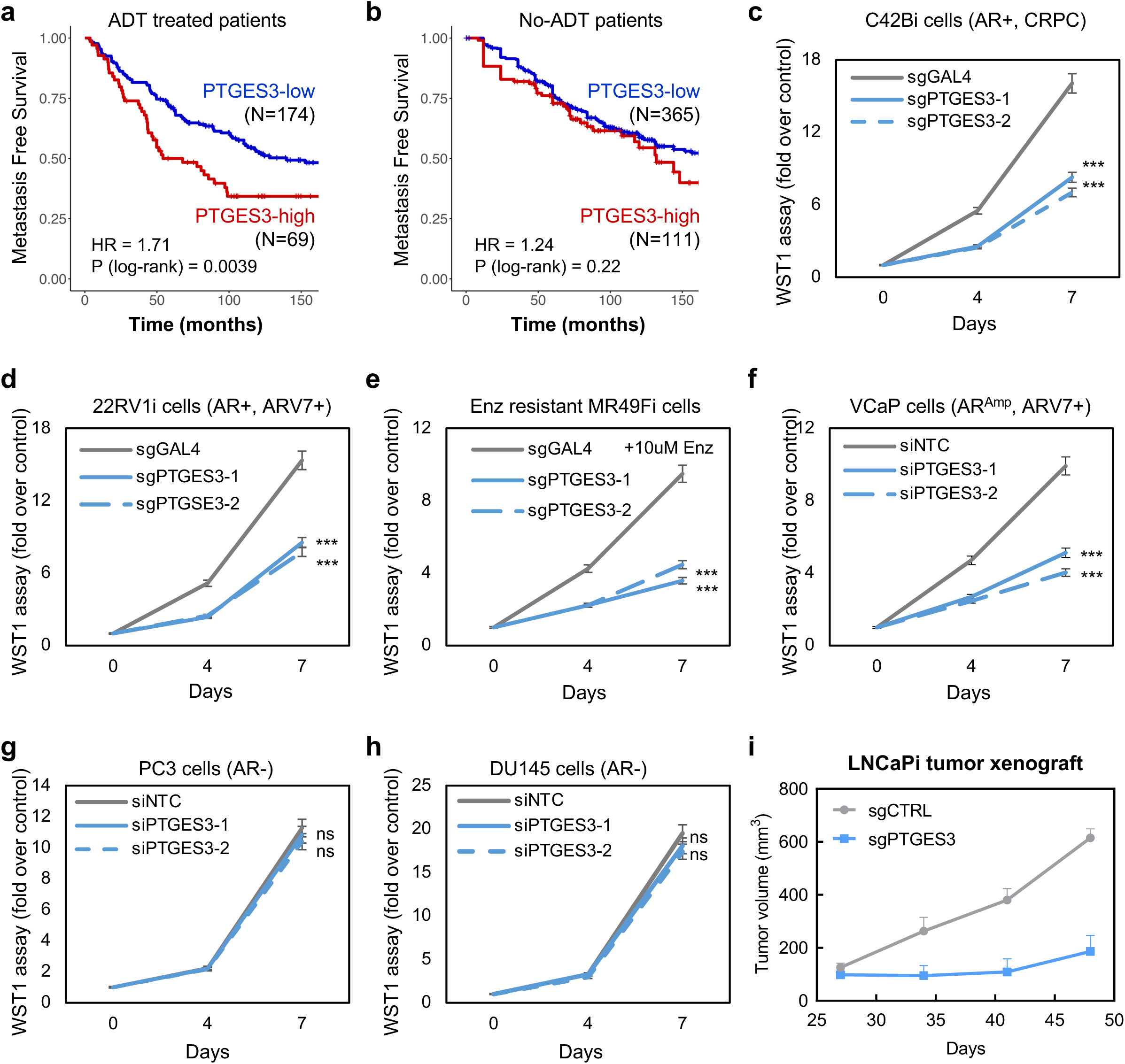
PTGES3 is a potential therapeutic target for AR-directed therapy resistant PCa. **a-b,** In a previously established cohort matched for 8 clinicopathologic prognostic variables^75^, high PTGES3 expression was associated with worse metastasis-free survival (MFS) in prostatectomy patients who received adjuvant ADT (HR:1.71, log-rank p-value=0.0039), but not in prostatectomy patients who did not receive adjuvant ADT (HR:1.24, log-rank p-value=0.22). **c-h,** C42Bi (**c**), 22RV1i (**d**), VCaP (**e**), or Enzalutamide (Enz) resistant cell line MR49Fi cultured with 5μM Enz (**e**), were infected with sgRNA targeting control GAL4 or PTGES3. VCaP (**f**), PC3 (**g**), or DU145 (**h**) cells were treated with siRNA targeting control (grey) or PTGES3 (blue). Cell viability was measured by WST1 and normalized over control (n = 3 as biological replicates; Mean ± SEM). Unpaired two-tailed t-test was used to determine statistical significance (ns = no significant difference; ***P < 0.001) **i,** Mice bearing LNCaP(TET-ON dCas9-KRAB) stably expressing sgRNA targeting GAL4 or sgPTGES3 were treated with doxycycline when tumors reached ∼200mm^3^. Average tumor volume was plotted and two-way ANOVA was used to measure statistical significance (n=8; * P<0.05).

With these promising clinical biomarker results, we further evaluated PTGES3 as a therapeutic target in multiple models of aggressive prostate cancer. We knocked down PTGES3 in multiple AR-dependent and AR-independent PCa cell lines and measured cell survival and proliferation. We found PTGES3 is required for cell proliferation in multiple AR-driven PCa models (C42B, 22RV1, VCaP, and MR49F cells) but not AR-independent PCa models (PC3 and DU145 cells) (**Fig.5c-h**). Notably, analysis of PTGES3 gene essentiality upon Cas9 knockout across 1070 diverse cancer cell lines present in the Cancer Dependency Map^50^ demonstrated that PTGES3 is not a commonly essential gene (**Extended data fig.10b**). Furthermore, CRISPRi experiments in other cancer cell types (AML, CML, pancreatic cancer and lung cancer cell lines^51–53^) demonstrated that unlike PSMA1, which is a positive control commonly essential gene, PTGES3 is not an essential gene upon knockdown (**Extended data fig.10c-f**). Additionally, PTGES3 repression led to a higher apoptotic population in AR-driven LNCaP cells (**Extended data fig.10g**). To test whether PTGES3 is required for AR-dependent prostate cancer cell growth or survival *in vivo*, we transduced an inducible CRISPRi LNCaP cell model (TET-ON dCas9-KRAB) with sgRNAs targeting PTGES3 or controls and transplanted cells into mice. At tumor onset we administered doxycycline to the mice to induce PTGES3 repression and tracked tumor size over time. We observed that repression of PTGES3 significantly delayed tumor growth *in vivo* relative to controls (**Fig.5i**). Analysis of AR protein levels in tumors demonstrated PTGES3 repression resulted in decreased AR protein abundance *in vivo* (**Extended data fig.10i**). Together, these results demonstrate that PTGES3 is required for AR levels and activity, is conditionally essential in diverse AR-driven PCa models but not non-AR-driven cancer cells and is associated with poor prognosis in PCa patients

## Discussion

We have created the first AR endogenous reporter and used this resource together with genome-scale CRISPRi screens to systematically identify networks of genes that sustain AR protein levels and thus AR’s oncogenic activity in prostate cancer. Our results point to unexpected biological activities that promote AR biology in aggressive prostate cancer. Specifically, we determined that PTGES3 affects AR levels and AR’s activity in the nucleus. Our results reveal that the PTGES3/AR interaction is required for mCRPC cell proliferation and viability *in vitro* and *in vivo*, including in models of patients with intrinsic and acquired AR directed therapy resistance. These findings emphasize the critical role of PTGES3 in the regulation of AR signaling and highlight its potential as a therapeutic target. Future therapeutic strategies targeting PTGES3 could be transformative for treating patients with AR-driven metastatic prostate cancer, especially those resistant to current FDA-approved Androgen Receptor Signaling Inhibitors (ARSIs). By targeting PTGES3, we may be able to overcome the significant challenge of drug resistance, addressing a major unmet medical need and ultimately improving survival outcomes for patients suffering from this lethal disease.

## METHODS

### Cell lines and reagents

Most cell lines were originally purchased from the American Type Culture Collection (ATCC) and were cultured per standard ATCC protocols. LNCaP, C42B, and 22RV1 were grown in the RPMI 1640 medium (Gibco) supplemented with 10% fetal bovine serum (FBS; Gibco), VCaP, DU145, PC3, and 293T cells in DMEM (ATCC) medium with 10% FBS. MR49F cells were a gift from the laboratory of A. Zoubeidi (Vancouver Prostate Center) and maintained in RPMI 1640 medium with 10% FBS containing 10uM Enzalutamide. Cells were grown in a humidified 5% CO2 incubator at 37°C. Cell line STR authentications were done at the UC Berkeley DNA Sequencing facility. All chemicals, unless otherwise stated, were purchased from Sigma Aldrich, Enamine, Combi-blocks or Astatech. Enzalutamide and Apalutamide were purchased from Selleckchem. ML-792 was purchased from MedKoo. AR PROTAC degrader ARD-61 was a gift from Dr. Shaomeng Wang’s lab in University of Michigan.

### CRISPRi screen

The genome-wide CRISPRi screens were performed following the Weissman lab protocol (weissmanlab.ucsf.edu) with some modifications. In brief, two individual clones of C42B^mNG2-AR^ cells were infected with a dCas9-KRAB construct (Addgene #46910). The genome-wide CRISPRi-V2 library (Addgene catalog #1000000091) virus was generated following the large scale lentivirus production protocol. An optimal MOI (<0.3) of the collected library virus to the C42Bi^mNG2-AR^ cells was determined by lentiviral titration.

For the flow based CRISPRi screen with the C42Bi^mNG2-AR^ cells, the sgRNA library virus was transfected at an average of 500x coverage after transduction (Day -7). Puromycin (5 µg/mL) selection for positively-transduced cells was performed 72 hours post transduction (Day -4). On Day 0, cells were fixed using 3% PFA, quenched by 30 mM glycine/PBS (pH 7.5), and collected for sorting with the BD FACSAria™ Fusion cell sorter. Cells were gated into the 25% of cells with the most mNG2 fluorescence intensity and 25% of cells with the least mNG2 fluorescence intensity. The screens were performed in two independent C42Bi^mNG2-AR^ clones. Genomic DNA was harvested with the QIAamp DNA FFPE Tissue Kit (cat# 56404). A standard genomic DNA PCR (22 cycles) was performed using NEB Next Ultra II Q5 master mix and primers containing TruSeq Indexes for NGS analysis. Sample libraries were prepared and sequenced on a HiSeq 4000. Screening data was analyzed using the established Weissman lab ScreenProcessing^35^ protocol (https://github.com/mhorlbeck/ScreenProcessing). sgRNA raw counts and phenotype scores calculated by the Weissman lab pipeline were included in **Supplementary table 2 and 3**. Gene phenotype scores calculated by the MAGeCK pipeline^54^ were included in **Supplementary table 4.**

### Plasmids, luciferase reporter assays, and siRNAs

The ORF of PTGES3 (addgene #108224) was cloned into pLVX-TetOne-Puro-C-Flag vector (gift from Dr. Minkyu Kim) using Gateway cloning (Invitrogen). Mutation constructs of PTGES3 were cloned with Q5® Site-Directed Mutagenesis (NEB). Guide RNAs expression vector targeting GAL4 or individual genes were cloned into pU6-sgRNA EF1Alpha-puro-T2A-mCherry or Lenti-sgRNA-blast (Addgene #104993). The ARE was cloned from ARR3tk plasmid (Addgene #132360) into pGL4 vector (Promega) to construct the ARE-luciferase reporter (ARE-luc). Luciferase activities were determined using the luciferin reagent (Promega) according to the manufacturer’s protocol. Transfection efficiency was normalized to Renilla luciferase activities. For siRNA knockdown experiments, cells were transfected with siRNA (**Supplementary table 1**) with Opti-MEM and Lipofectamine RNAiMAX (Invitrogen) at the final concentration of 50 nM, and incubated for at least 48 hours. Non-targeting control siRNA was used as the control.

### STRING and Gene Set Enrichment Analysis (GSEA)

For protein network analysis, the top 51 positive AR regulator hits from the screen were submitted to STRING (V11.5)^37^. Genes without gene interactions among any of the top 51 hits were removed. Physical subnetwork was chosen. Connections were based on confidence. Line thickness indicates the strength of data support. Genes were clustered by unsupervised Markov Cluster Algorithm (MCL clustering, inflation parameter=3). For CRISPRi screening hits, gene set enrichment analyses were performed using the fgsea R package (V1.16.0), using all genes ranked by the screening score (log10 p-value x phenotype z) for the molecular signatures database (MSigDB) Cancer Hallmarks collection and the curated Wikpathways collection. Significantly up or down regulated pathways (adjusted *p* < 0.05) are shown. For RNA-seq, gene set enrichment analyses were performed using the pre-ranked method implemented in the fgsea R package (10.18129/B9.bioc.GSEABase), and Hallmark gene sets were downloaded from the molecular signatures database (MSigDB), genes were ranked by the Wald-statistics from DESeq2.

### RNA-extraction and qPCR

RNA was extracted from the cells as per manufacturer’s protocol using the Zymo Quick-RNA extraction kit (Cat# R1054). cDNA was prepared using SuperScript III First-Strand Synthesis System (Cat# 18080). The mRNA expression of the genes was measured using primers (**Supplementary table 1)** and SYBR Green Real-Time PCR Master Mixes (Thermo) in the QuantStudio Flex Real-Time PCR system.

### Western blot, far western blot and Immunoprecipitation

Protein from the cells was extracted with either RIPA buffer with protease inhibitor (Thermo, Cat#78430) or a cell fractionation kit (Cell signaling #9038S). Western blots were performed as described previously^55^. Antibody information were listed in the **Supplementary table 5.** Far western blot analysis was performed as detailed^56^. In brief, 293T cells were transfected with pCMV-Flag-AR (gift from Dr. Paul Rennie in Vancouver Prostate Center) using Lipofectamine 3000 (Invitrogen). AR was immuno-affinity purified with an anti-FLAG and resolved by SDS-PAGE, transferred to PVDF membrane, and denatured and renatured. After blocking, membranes were incubated with probes then blotted for PTGES3 antibody. For immunoprecipitation, protein lysates were incubated with primary antibody overnight. Immunoprecipitation experiments were performed as previous described^55^. Briefly, pre-cleared cell lysates were incubated with primary antibody and the associated proteins were immunoblotted by indicated antibodies. Each experiment was performed twice to determine reproducibility, representative images were shown.

### Mass Spectrometry

Approximately 10 million cells were lyesed using Lyse buffer from the PreOmics iST Kit (PreOmics GmbH; Martinsried, Germany). The samples were reduced and alkylated at 90 °C for 10 minutes and continuously shaken at 1000 RPM. 100 μg clarified protein samples were enzymatically digested using Trypsin/Lys-C mix at 37 °C for 90 minutes and peptides were desalted following the PreOmics iST kit manufacturer’s protocol and dried down in a speedvac overnight at room temperature (CentriVap, Labconco). The peptide samples were resuspended in Solvent A (2% ACN, 0.1% FA) and quantified using the Pierce Quantitative Peptide Assay (Thermo Scientific). A total of 1 μg of peptides from each sample were loaded on to an EASY-Spray nanocolumn (Thermo Fisher Scientific, ES900) installed on Dionex Ultimate 3000 NanoRSLC coupled with Q-Exactive Plus mass spectrometer (Thermo Fisher Scientific). Peptides were separated over a 95-minute gradient of ACN ranging from 2% to 30% ACN followed by a quick ramp up to 80% ACN. MS/MS scans were performed over mass range of m/z 350-1500 with resolution of 17,500. The isolation window was set to 4.0 m/z and the charge state was set to 2.

The MS/MS raw data (.raw files) were processed using MSFragger within FragPipe^57^ with default Label Free Quantification (LFQ) settings and searched against the human uniport database. The contaminant and decoy protein sequences were added to the search database via Fragpipe. The computational tools PeptideProphet and ProteinProphet were used for statistical validation of results and subsequent mapping of the peptides to the proteins respectively with 1% FDR. The LFQ intensities were considered for secondary data analysis. Features having missing values of more than 50% were eliminated, and KNN was used to impute the missing values. The data were log transformed, normalized and subjected to Welch’s t-tests (**Supplemental table 6**). Proteins with p-values less than 0.05 were used to identify the differentially expressed proteins. Statistical analysis and data visualization were performed in R studio and MetaboAnalyst 5.0^58^.

### Immunohistochemistry (IHC) and staining score evaluation

The prostate tumor microarray (TMA) with correlated clinical information from 120 clinically localized prostate cancer patients was constructed by the University of Michigan Center for Translational Pathology as previously reported^59^. IHC was performed as previously described^60^. Briefly, paraffin embedded TMA slides were deparaffinized and rehydrated following standard protocols. Antigen retrieval was carried out with HIER EDTA buffer. Endogenous peroxidase activity was blocked using 1% hydrogen peroxide. Slides were probed overnight at 4°C with anti-PTGES3 (1:20, Sigma HPA038672), washed, incubated with biotinylated secondary anti-rabbit antibody (Jackson #111-065-144, 1:2000), followed by incubation with streptavidin-HRP (Invitrogen #SA10001, 1: 250). Staining was visualized using DAB developing kit (Vector Laboratories) and nuclei were counterstained with hematoxylin (Vector Laboratories). The stained TMA slide was scanned by a Leica Aperio AT2 scanner and digital images were evaluated independently by two pathologists, B.A.S. and J.Z. IHC scores of nuclear PTGES3 were calculated by IHC signal intensity (0-1 as weak and strong as 2-3).

### Chromatin immunoprecipitation (ChIP) qPCR and Dual X ChIP qPCR

ChIP and Dual X ChIP assays were performed as previously described^55,61^. In brief, 1x10^7^ cells were collected, washed, and cross-linked with 1% formaldehyde in RT for 10 mins. For Dual X ChIP crosslinking, cells were treated with 1.5 mM Ethylene glycol bis (EGS in PBS, Sigma) for 30 mins, subsequently crosslinked with 1% formaldehyde in RT for 10 mins, and quenched with 1.25 M Glycine for 5 mins. Genomic DNA extraction and ChIP assay were then performed using Bioruptor Pico sonication (Diagenode) and HighCell# ChIP kit (Diagenode, C01010061). Bound DNA was quantified by quantitative PCR (SYBR Green master mix, Invitrogen) using the primer sets listed in **Supplementary table 1**. The quantitative PCR results are presented as fold enrichment of PCR amplification over control IgG antibody and normalized based on the total input (non-precipitated chromatin). Primers for the GAPDH promoter were used as a negative control.

### RNA seq

RNA-Seq sample preparation: LNCaPi cells were viral transduced with sgRNA targeting PTGES3 or control sgRNA for 3 days and selected with puromycin (5 µg/mL) to purify the population for 3 days. Post-selection RNA was extracted from the cells as described above to generate Illumina compatible libraries and perform qPCR to determine the sgRNA knockdown efficiency. The QuantSeq 3’mRNA-Seq library prep kit FWD for Illumina (Lexogen, Cat# 015.24) was used to prepare the library as per the manufacturer’s protocol. Quality control was performed by using the Agilent Bioanalyzer 2100 system and the samples were sequenced by an Illumina HiSeq 4000. The entire RNA-seq experiment was done in two biological replicates. The RNA-seq single-end fastq data generated by Illumina HiSeq 4000 sequencing system were first trimmed to remove adapter sequences using Cutadapt v2.6 with the “-q 10 -m 20” option^62^. After adapter trimming, FASTQC v0.11.8 was used to evaluate the sequence trimming as well as overall sequence quality. Using the splice-aware aligner STAR (2.7.1a, ^63^), RNA-seq reads were aligned onto the Human reference genome build GRCh38decoy using the “--outSAMtype BAM SortedByCoordinate --outSAMunmapped Within -- outSAMmapqUnique 50 --sjdbOverhang 65 --chimSegmentMin 12 --twopassMode Basic” option and exon-exon junctions, with Human gene model annotation from GENCODE v30. Gene expression quantification of uniquely mapping reads was performed using the “featurecount” function within Rsubread R-package^64^ with “GTF.featureType=“exon”, GTF.attrType=“gene_id”, useMetaFeatures=TRUE, allowMultiOverlap=FALSE, countMultiMappingReads=FALSE, isLongRead=FALSE, ignoreDup=FALSE, strandSpecific=0, juncCounts=TRUE, genome=NULL, isPairedEnd=FALSE, requireBothEndsMapped=FALSE, checkFragLength=FALSE, countChimericFragments=TRUE, autosort=TRUE” option. Cross-sample normalization of expression values and differential expression analysis between the PTGES3 knockdown and control was done using DESeq2 R-package^65^. Benjamini-Hochberg corrected p-value < 0.05 and log2 foldchange > 1 or < -1 were considered statistically significant.

### ATAC seq

We performed ATAC-Seq on LNCaP cells following knockdown of PTGES3 or control. The experiment was carried out as described in the published method papers by Buenrostro, et al.^66^ and Corces, et al.^67^ with the following modifications. Cells were resuspended in buffer (Illumina Cat#20034198), incubated on ice for 10 min, and lysed using a dounce homogenizer. 50,000 nuclei were incubated with 25 µL 2x TD Buffer and 1.25 µL Transposase (Illumina Tagment Enzyme/Buffer Cat# 20034210) shaking at 300 rpm at 37°C for 30 min. Zymo DNA Clean and Concentrator 5 kit (Cat# D4014) was then used to purify DNA. Transposed DNA was amplified using PCR master mix and indexes from Nextera DNA Library Prep kit (Cat# 15028211) for 5 cycles and then assessed using qPCR. Final cleanup was performed using 1.8x AMPure XP beads (Cat# A63881) and libraries quantified using the DNA High Sensitivity Agilent 2100 Bioanalyzer System. Samples were sequenced at the UCSF Core Facility on a HiSeq4000 with PE100 libraries. The entire experimental setup was performed in two technical replicates.

*ATAC-seq data processing*: The ATAC-seq paired-end fastq files were first trimmed to remove Illumina Nextera adapter sequence using Cutadapt v2.6^68^ with the “-q 10 -m 20” option. After adapter trimming, FASTQC v0.11.8^69^ was used to evaluate the sequence trimming as well as overall sequence quality. Bowtie2 version 2.3.5.1^70^ was then used to align the ATAC-seq reads against the Human reference genome build GRCh38decoy using the “--very-sensitive” option. The uniquely mapped reads were obtained in SAM format. Samtools version 1.9^71^ was used to convert SAM to BAM file as well as sort the BAM file. Picard (https://broadinstitute.github.io/picard/) was then used to flag duplicate reads using the MarkDuplicates tool using “REMOVE_DUPLICATES=true” option. The resulting BAM file reads position were then corrected by a constant offset to the read start (“+” stranded +4 bp, “-” stranded -5 bp) using deepTools2 v3.3.2^72^ with “alignmentSieve --ATACshift” option. This resulted in the final aligned, de-duplicated BAM file that was used in all downstream analyses. ATAC-seq peak calling was performed using MACS2 v2.2.5^73^ to obtain narrow peaks with “callpeak -f BAMPE -g hs –qvalue 0.05 --nomodel -B --keep-dup all --call-summits” option. The resulting peaks that map to the mitochondrial genome or genomic regions listed in the ENCODE hg38 blacklist (https://www.encodeproject.org/annotations/ENCSR636HFF/) or peaks that extend beyond the ends of chromosomes were filtered out. Non-overlapping unique ATAC-seq narrow peak regions were obtained from all samples analyzed. Only those non-overlapping unique peak regions present in at least two samples were considered for further analysis. Sequencing reads mapped to these non-overlapping unique regions were counted using “featurecount” function within Rsubread^63^ R-package with “isPairedEnd=TRUE, countMultiMappingReads=FALSE, maxFragLength=100, autosort=TRUE” option. Further library-size normalization of the feature counts and differential open chromatin regions between sgPTGES3 and sgGAL4 were obtained using DESeq2^65^ R-package. Only those peak regions with Benjamini-Hochberg corrected pvalue < 0.05 and log2 foldchange > 1 or < -1 were considered statistically significant. The ATAC-seq peaks were annotated using ChIPseeker^74^ R-package based on hg38 GENCODE v30 annotations.

Over-representation Analysis: The nearest gene to the ATAC-seq peaks that mapped within the promoter regions of the corresponding gene were considered for this analysis. These genes were tested for enrichment against Hallmark genesets in Molecular Signature Database (MSigDB) v7.0. A hypergeometric test-based over-representation analysis was used for this purpose. A cut-off threshold of false discovery rate (FDR) ≤ 0.01 was used to obtain the significantly enriched genesets.

Transcription Factor (TF) binding analysis: The differentially accessible ATAC-seq peaks that mapped to promoter and intergenic regions were used for TF binding analysis. The MEME tool^75^ was used to discover TF binding motifs de-novo. These potential TF binding motifs were annotated for known TF motif from the JASPAR^76^ database using the TomTom tool within the MEME tool suite.

### Clinical cohorts’ analysis

Publicly available gene expression data from a matched cohort established previously^77^ or mCRPC was downloaded from cBioportal (2021-05-20)^78–80^. Samples were grouped based on expression levels above or below the 25th percentile for PTGES3 separately for poly-A or capture-based RNA-seq, and the capture-based results were used if a sample had sequencing data available from both methods. Differences in survival between groups were visualized using the Kaplan-Meier method and a logrank-test was used to test for differences in survival, using the survival and survminer R packages^81,82^. Hazard ratios were calculated using Cox proportional hazards regression.

### Tumor Models

Doxycycline inducible LNCaPi cells established previously^83^ were lentiviral transfected with sgRNA targeting PTGES3 and control-sgRNA. Post puromycin selection, cells were counted, and the cell suspension was mixed with Matrigel (Corning #354230, 1:1). 2x10^6^ cells were injected subcutaneously on each flank of the mice. Male NOD-SCID-Gamma (NSG) mice (6-8 weeks) were used. Once the tumors were palpable, the mice were randomly categorized in two groups receiving either doxycycline diet (Bio-Serv, Cat#S3888) or control diet. Tumors were measured using digital calipers. Tumor volume was calculated using the equation: Volume= length x width^2^ x 0.52, where the length represents the longer axis. Average tumor volume was plotted and Two-way ANOVA was used to measure statistical significance denoted by asterisk (*). *p < 0.05, **p < 0.01, *** p < 0.001. The mice were humanely euthanized following appropriate UCSF’s laboratory animal resource center (LARC) protocol once the tumor volume reached 1000mm^3^. All animal experiments conducted were reviewed and approved by UCSF IACUC.

### WST-1, clonogenic, and IncuCyte assays

Cell viability assays were performed as previously described^55^ using the cell proliferation reagent WST-1 (Sigma) according to the manufacturer protocol. Clonogenic assays were performed as previous described^83^. In brief, 1000 cells were seeded in a 6-well plate and incubate with indicate treatments for 10 days. The cell colonies were then washed with PBS and fixed/stained with 25% methanol plus crystal violet (0.05% w/v). Images were scanned and analyzed using a GelCount (Oxford Optronix). Average number of colonies formed were counted. For IncuCyte experiment, cells labeled with Nuclight red were seeded to 96 well plates. The time relapse images were captured and analyzed with by the Incucyte® S3 live-cell analysis system. Each experimental setup was performed in three biological replicates and done at three times to determine the reproducibility of the data.

### Statistical analysis

Spearman’s correlation was used to determine statistical significance for all the correlation plots. For gene expression and correlation, the Wilcoxon rank-sum test was used to test for differences between two groups, unless otherwise stated. Unpaired t-test were used to determine statistical analysis for the column plots, denoted by asterisk (*). *p < 0.05, **p < 0.01, ***p < 0.001. Two-way ANOVA was used to determine statistical significance in the *in vivo* data. In RNA-Seq data, the Benjamini-Hochberg test was performed. Corrected p-value < 0.05 and log2 foldchange > 0.5 or < 0.5 were considered statistically significant. In ATAC-Seq data peak regions with Benjamini-Hochberg corrected p-value < 0.05 and log2 foldchange > 0.5 or < 0.5 were considered statistically significant.

### Data availability

RNA seq and ATAC seq data is available at GEO (GSE180036). ChIP-seq data used is available at GEO (GSE120741).

## ACKNOWLEDGEMENTS

We thank all members of the Feng lab as well as the Gilbert lab for helpful suggestions and technical advice. We thank Dr. Manuel Leonetti for helpful advice on fluorescently engineered cell lines. We thank Dr. Arul Chinnaiyan and Dr. Shaomeng Wang for sharing the AR degrader. We thank Dr. Amina Zoubeidi for sharing the enzalutamide resistant cell line. We thank Dr. Alan Ashworth, Dr. Jason Gestwicki, and Dr. Charlotte Bevan for helpful advice. H.L. was supported by the Prostate Cancer Foundation Young Investigator Award and the UCSF Prostate Cancer Program 2021 Pilot Research Awards. J.E.M. was supported by National Institute of Health (NIH) 1F32CA236347-01. M.S. was supported by the Swedish Research Council (Vetenskapsrådet) with grant number 2018-00382, the Swedish Society of Medicine (Svenska Läkaresällskapet) and the Prostate Cancer Foundation Young Investigator Award. L.C. was supported by the Prostate Cancer Foundation Young Investigator Award and the Department of Defense Prostate Cancer Research Program Early Investigator Research Award. J.C. was funded by a Prostate Cancer Foundation Young Investigator Award and DOD award W81XWH-20-1-0136. J.T.H. was funded by a Prostate Cancer Foundation Young Investigator Award. B. H. was supported by National Institutes of Health (NIH) R01GM124334 and R01GM131641. B. H. is a Chan Zuckerberg Biohub Investigator. E.J.S was supported by a Stand Up To Cancer-Prostate Cancer 553 Foundation Prostate Cancer Dream Team Award (SU2C-AACR-DT0812). D.A.Q. was funded by a Prostate Cancer Foundation Young Investigator Award and a BRCA Foundation Young Investigator Award. K.M.S. is supported by the Howard Hughes Medical Institute, Samuel Waxman Cancer Research Foundation, National Institute of Health (NIH) 1R01CA221969-01 and National Institute of Health (NIH) 1R01CA244550. L.A.G. was supported by K99/R00 CA204602 and DP2 CA239597, as well as the Goldberg-Benioff Endowed Professorship in Prostate Cancer Translational Biology. F.Y.F. was supported by National Institutes of Health (NIH)/National Cancer Institute (NCI) 1R01CA230516-01. F.Y.F. was supported by NIH/NCI 1R01CA227025 and Prostate Cancer Foundation (PCF) 17CHAL06. F.Y.F. was supported by NIH P50CA186786. This project is funded by Prostate Cancer Foundation Challenge award to L.A.G., F.Y.F, and K.M.S. Additional funding was provided by a UCSF Benioff Initiative for Prostate Cancer Research award.

## AUTHOR CONTRIBUTIONS

H.L., J.E.M., L.A.G. and F.Y.F. conceived and designed the study.

H.L., J.E.M, B.F., S.F., M.S., L.C., M.C., R.D., E.A.E., M.A., H.S., L.A.G., and F.Y.F. acquired the data.

H.L., J.E.M, B.F., S.F., M.S., R.S., M.Z., L.C., M.C., J.C., R.D., E.A.E., A.W., J.Z., A.M., J.T.H., M.A., W.S.C, B.A.S., J.S., B.H., E.J.S., D.A.Q., K.M.S., L.A.G., and F.Y.F. analyzed or interpreted the data.

All authors drafted the article or revised it critically for important intellectual content. All authors approved the final version of the manuscript.

## TABLES

**Supplemental Table 1** Synthetic oligos

**Supplemental Table 2** clone 1 screen data

**Supplemental Table 3** clone 2 screen data

**Supplemental Table 4** screen data processed by MAGeCK

**Supplemental Table 5** Antibody list

**Supplemental Table 6** Mass spectrometry data

---

**Extended data figure 1.**
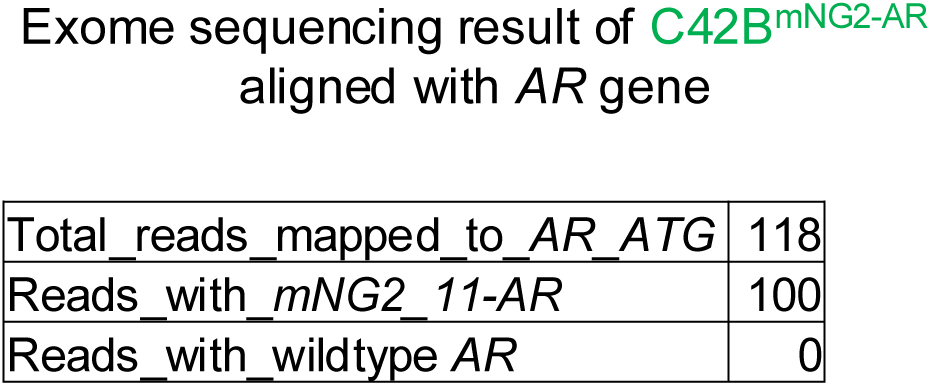
Exome sequencing results of AR reporter cell line. Exome sequencing results of C42B^mNG2-AR^ showed a homogenous knock-in of mNG2.

**Extended data figure 2.**
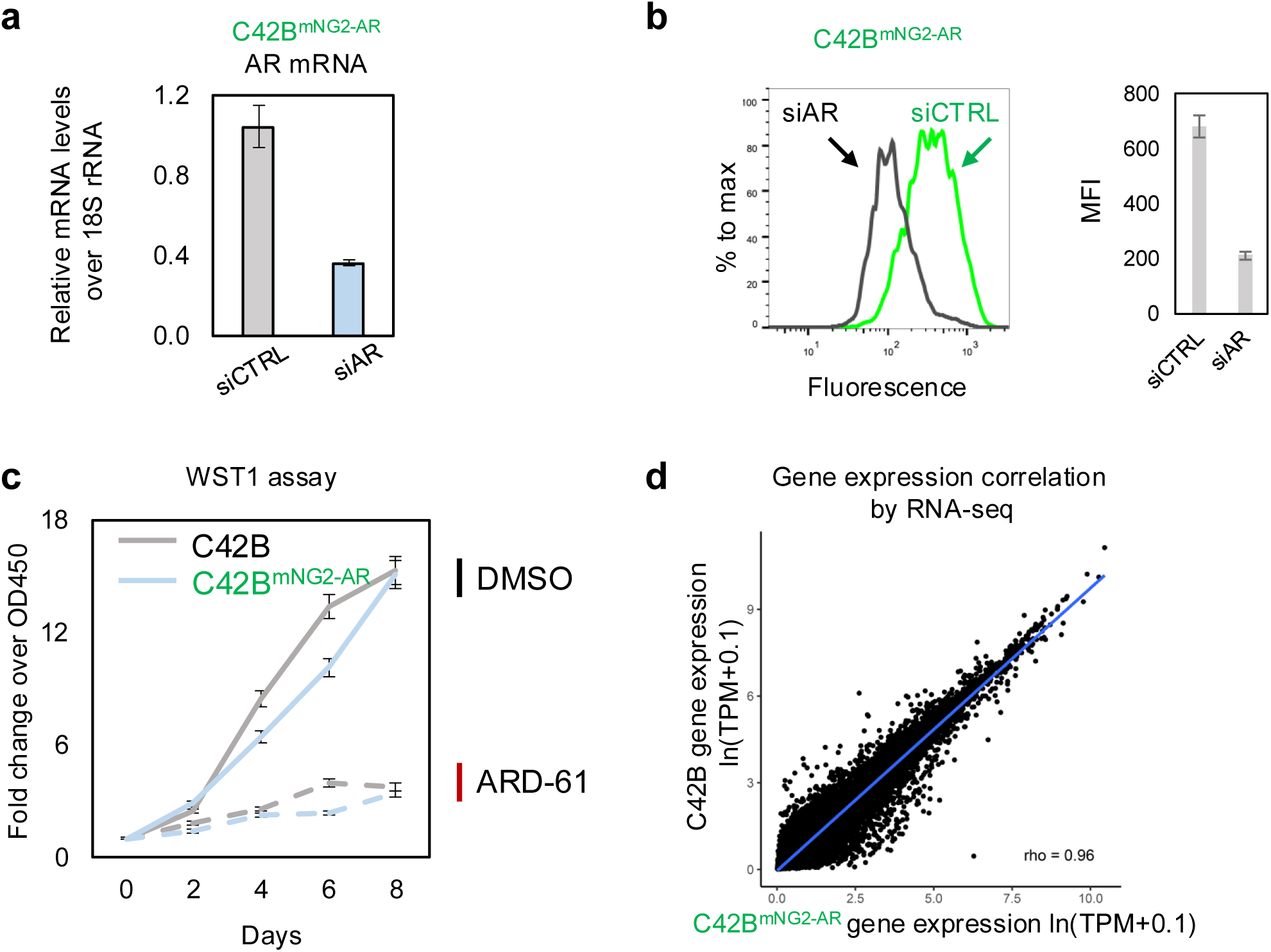
Phenotypical characterization of C42B^mNG2-AR^ cell lines. **a,** C42B^mNG2-AR^ cells were treated with siRNA targeting control (grey) or AR (blue). RNA was collected, AR mRNA levels over 18s rRNA were measured by real-time PCR then normalized to siCTRL (n = 3 as biological replicates; Mean ± SEM). **b,** C42B^mNG2-AR^ cells were treated with siRNA targeting control (green) or AR (black). Cells were collected for flow cytometry, mNG2-AR levels were indicated by the fluorescence intensity. Median Fluorescence Intensity (MFI) was calculated (n = 3; Mean ± SEM). **c,** C42B (grey) and C42B^mNG2-AR^ cells (blue) were treated with DMSO (solid line) or 50nM ARD-61 (dashed line). Cell viability was measured by WST1 and normalized over control (n = 3 as biological replicates; Mean ± SEM). **d,** Total RNA from C42B and C42B^mNG2-AR^ cells were collected for RNA-seq. Gene expression values were calculated as ln(TPM+0.1). Pearson correlation was calculated comparing all genes (n=19127) between the two cell lines.

**Extended data figure 3.**
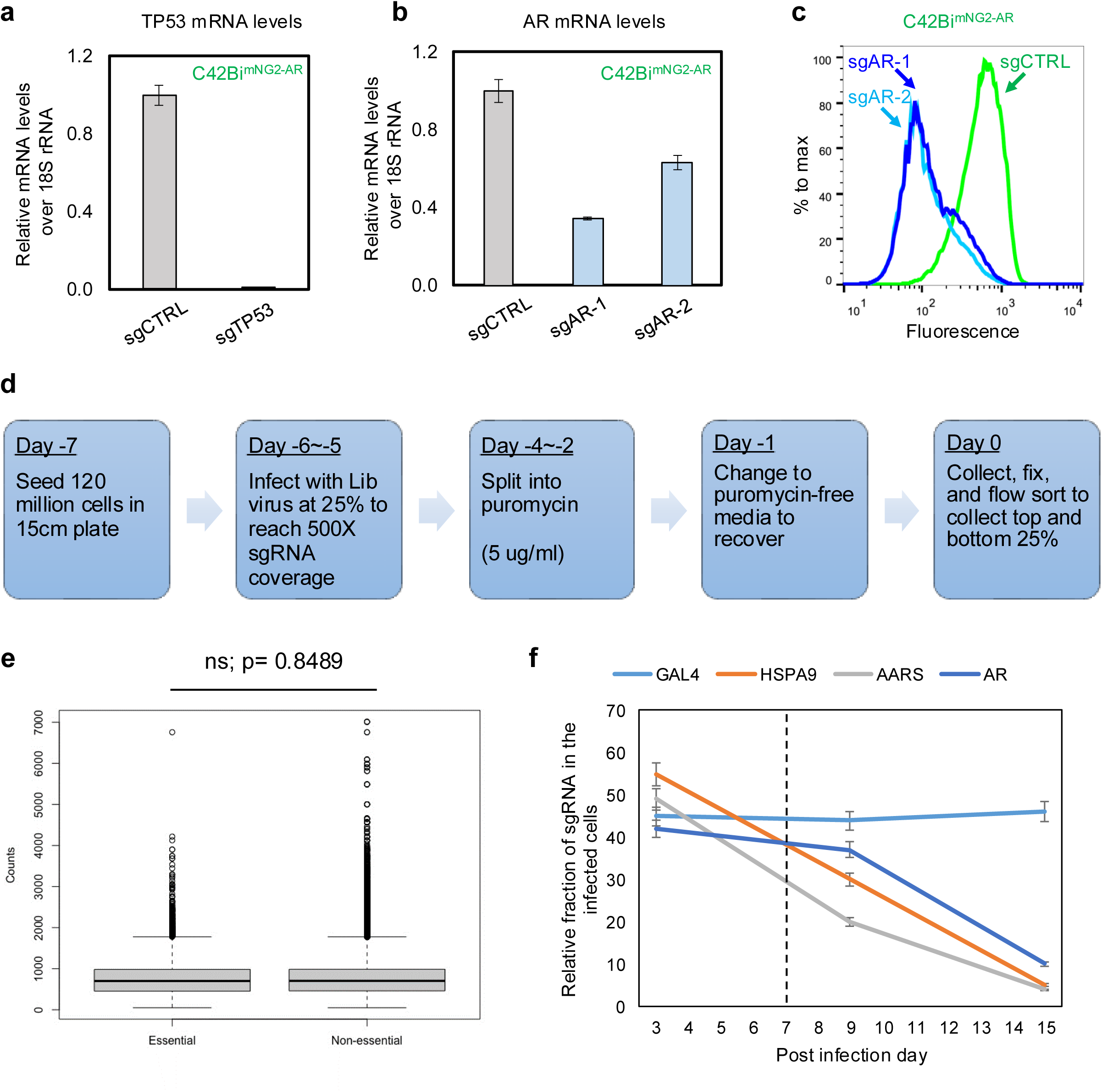
Validation of the CRISPRi activity of C42Bi^mNG2-AR^. **a-b,** C42Bi^mNG2-AR^ cells were infected with sgRNAs targeting GAL4, P53 (**a**) or AR (**b**). RNA was collected; P53 or AR mRNA levels over 18s rRNA were measured by real-time PCR (n = 3 as biological replicates; Mean ± SEM). **c**, C42Bi^mNG2-AR^ cells were infected with sgRNAs targeting control (green) or AR (blue). Cells were collected for flow cytometry; mNG2-AR levels were indicated by the fluorescence intensity (n = 3 as biological replicates; Mean ± SEM). **d**, Flow chart illustrates the details of the flow base CRISPRi AR reporter screen. **e**, Counts of essential and non-essential sgRNAs at post infection day 7 (T0) were shown. There was no significant difference between the two groups. **f**, A competitive growth-based assay of the C42Bi^mNG2-AR^ cells infected with sgRNA (mCherry+) targeting gene of interest or control. The relative fraction of sgRNA in the cell population was measured by flow cytometer starting from day 3 post-infection.

**Extended data figure 4.**
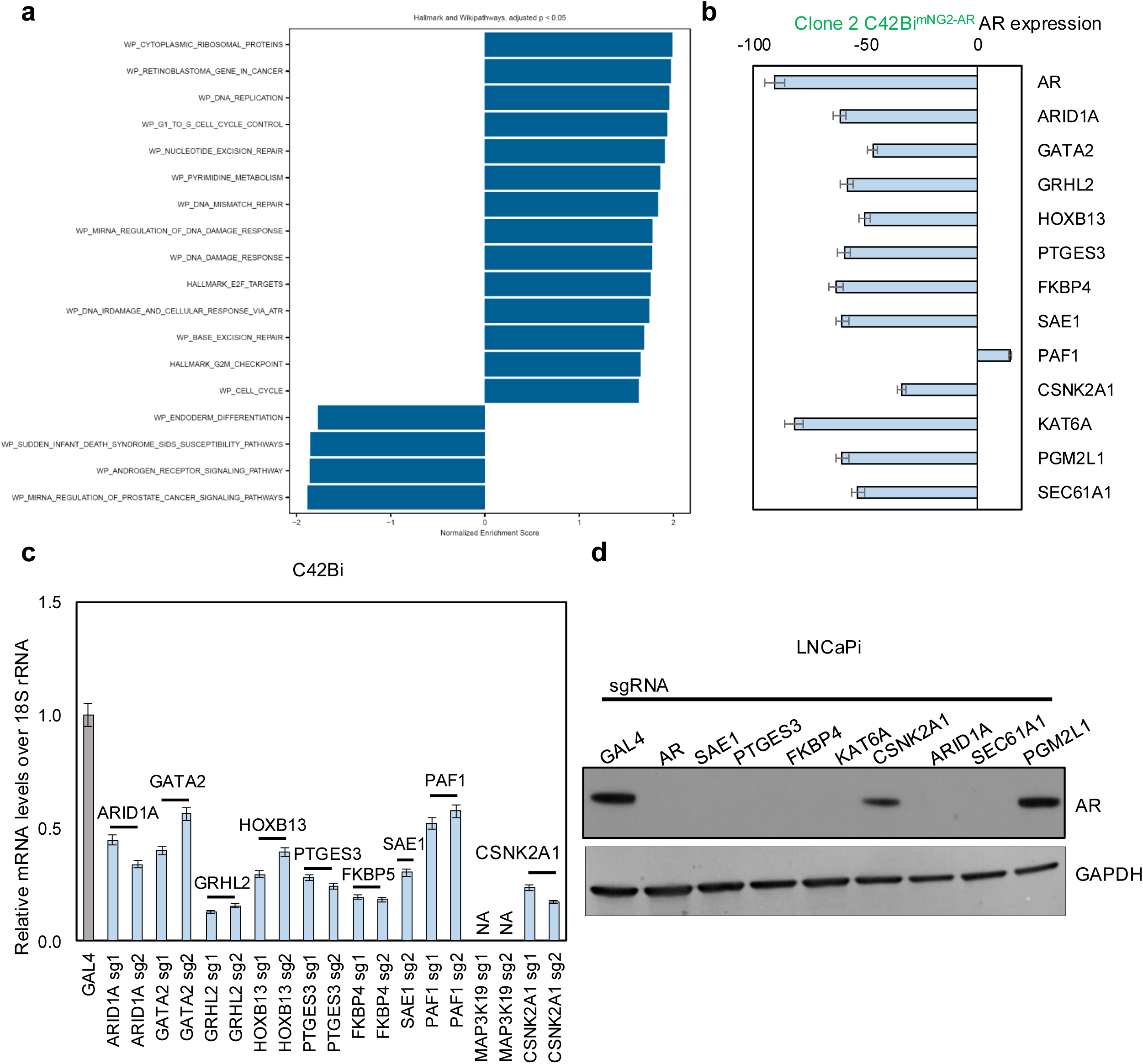
Extended validation of hits from the genome-wide CRISPRi screening. **a,** Screen hits were analyzed with Fast Gene Set Enrichment Analysis with fgsea multilevel function (fGSEA V1.16.0). **b,** Clone 2 C42Bi^mNG2-AR^ were infected with sgRNAs targeting control or indicated genes. After puromycin selection, Median Fluorescence Intensity (MFI) of mNG2-AR fluorescence from the treated cells was measured by flow cytometry. The percentage of AR expression was calculated as AR expression%=(MFI_C42B_-MFI_sg **Individual gene**_)/(MFI_C42B_-MFI_sgGAL4_)*100 (n = 3 as biological replicates; Mean ± SEM). **c,** C42Bi were infected with sgRNAs targeting GAL4 control or indicated genes. mRNA levels of the indicated genes over 18s rRNA were measured by real-time PCR (n = 3 as biological replicates; Mean ± SEM). **d,** LNCaPi cells were infected with indicated individual sgRNAs. Cell lysates were collected after puromycin selection. AR and GAPDH levels were detected by western blotting. Each western blot experiment was performed twice to determine reproducibility. **e,** C42B cells were treated with 100nM ARD-61 (ARi), 5μM ML792 (SAEi), siRNA targeting PTGES3, or 10μM Slimitasertib (CSNK2A1i) for 24h. Cell lysates were collected. AR and GAPDH levels were detected by western blotting.

**Extended data figure 5.**
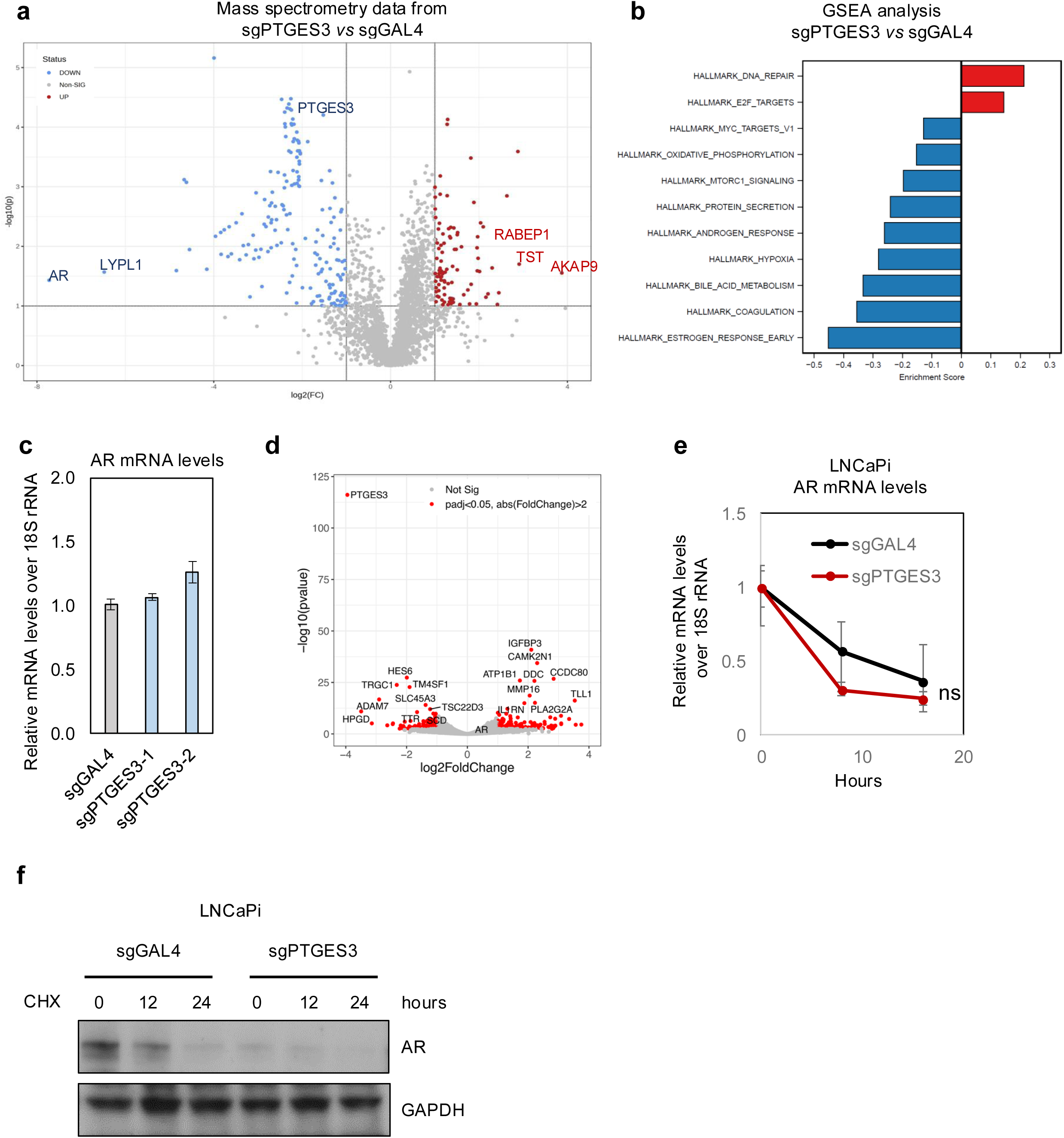
PTGES3 does not affect AR mRNA levels and is not required for cell proliferation or survival in most non-prostate cancer or AR independent cells. **a**, LNCaPi cells were infected with sgRNA targeting GAL4 (control) or PTGES3. Cells were collected for label free shotgun mass spectrometry after puromycin selection (n=3). Volcano plot showing the differentially expressed proteins in sgPTGES3 vs sgGAL4 groups. **b**, Gene Set Enrichment Analysis (GSEA) was performed. **c,** LNCaPi were infected with sgRNAs targeting GAL4 control or PTGES3. AR mRNA levels over 18s rRNA were measured by real-time PCR (n = 3 as biological replicates; Mean ± SEM). **d,** LNCaPi were infected with sgRNA targeting control or PTGES3. Total RNA was collected for RNA-seq. Differential expressed (DE) genes between the two groups were highlighted in red in the volcano plot; top ranking DE genes were labeled. AR gene is not differentially expressed when PTGES3 is knocked down. **e.** LNCaPi cells infected with sgGAL4 or sgPTGES3 were treated with 1uM Actinmycin D for 0, 8 or 16 hours. RNA was collected to measure the AR mRNA levels using real-time PCR (n = 3 as biological replicates; Mean ± SEM; ns indicates no significant difference by two-way ANOVA analysis). **f.** LNCaPi cells infected with sgGAL4 or sgPTGES3 were treated with protein synthesis inhibitor Cycloheximide (CHX, 5μM) for 0, 12, 24 hours. AR protein levels were detected by western blotting.

**Extended data figure 6.**
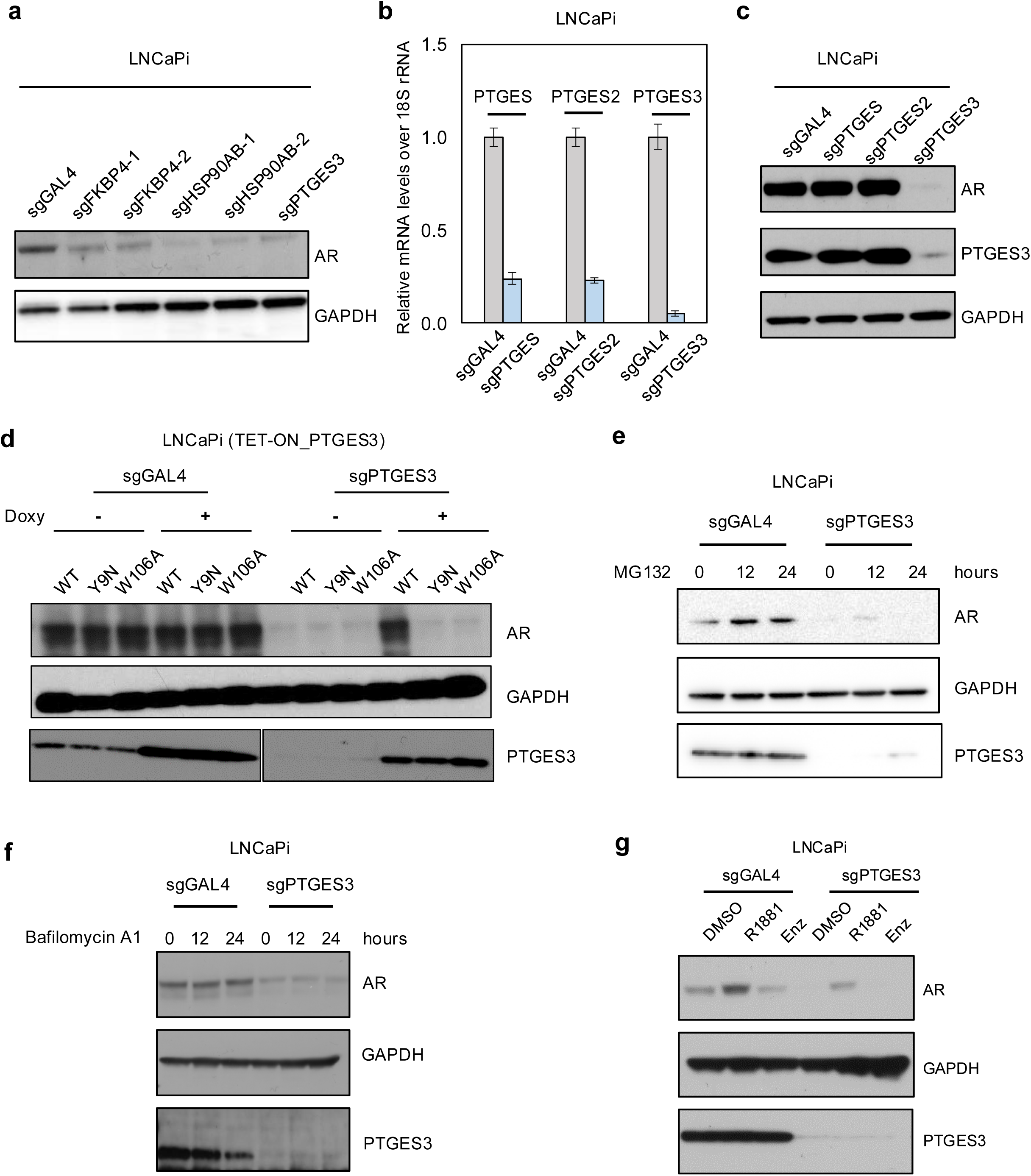
Dual functions of PTGES3 potentially regulating AR protein levels. **a,** LNCaPi were infected with sgRNA targeting GAL4, FKBP4, HSP90 or PTGES3. Cell lysates were collected. AR and GAPDH levels were detected by western blotting. **b-c,** LNCaPi were infected with sgRNA targeting GAL4, PTGES, PTGES2 or PTGES3. RNA was collected, indicated genes mRNA levels over 18s rRNA were measured by real-time PCR (**b;** n = 3 as biological replicates; Mean ± SEM). Cell lysates were collected. AR and GAPDH levels were detected by western blotting (**c**). Each western blot experiment was performed twice to determine reproducibility. **d,** LNCaPi cells were infected with sgGAL4-Blast or sgPTGES3-Blast, then treated with DMSO or 100ng/ml doxycycline to overexpress the PTGES3 WT or Y9N or W106A mutant proteins. AR and GAPDH levels were detected by western blotting. **e,** LNCaPi cells infected with sgGAL4 or sgPTGES3 were treated with proteasome inhibitor 5μM MG132 for 0, 12, 24 hours. AR protein levels were detected by western blotting. **f,** LNCaPi cells infected with sgGAL4 or sgPTGES3 were treated with lysosomal inhibitor 1μM Bafilomycin A1 for 0, 12, 24 hours. AR protein levels were detected by western blotting. **g,** LNCaPi cells infected with sgGAL4 or sgPTGES3 were treated DMSO, 10nM R1881 (androgen), or 10μM Enzalutamide (Enz, anti-androgen). AR protein levels were detected by western blotting.

**Extended data figure 7.**
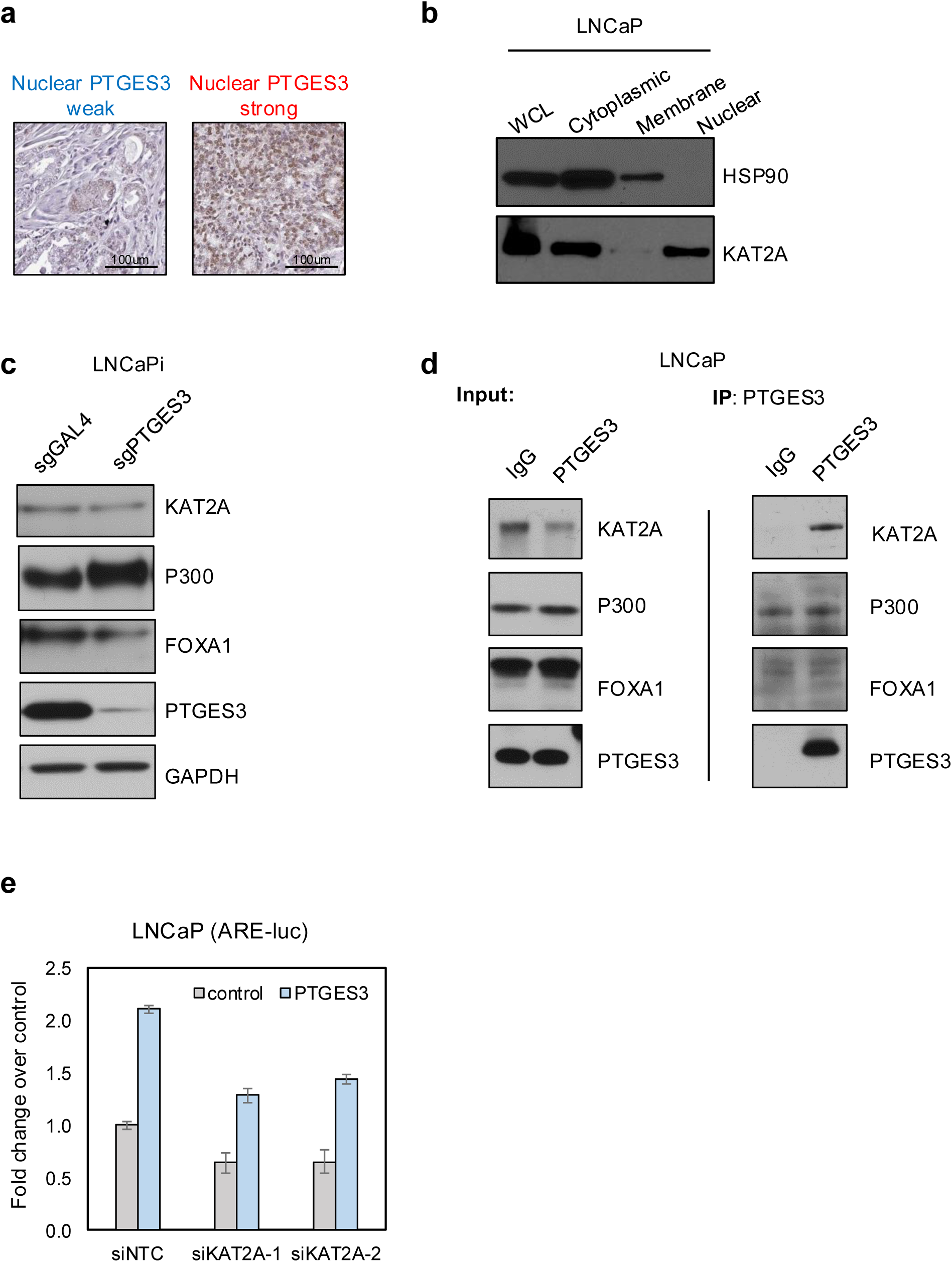
PTGES3 has a nuclear HSP90-independent localization and potentially interacts with histone acetyltransferase KAT2A. **a,** Representative immunostainings against PTGES3 in PCa patient samples. **b,** Cell fractions from LNCaP cells were immunoblotted with indicated antibodies. **c,** LNCaPi cells were infected with sgGAL4 or sgPTGES3. KAT2A, P300, FOXA1, PTGES3, and GAPDH protein levels were detected by western blotting. **d,** IP experiments were performed with LNCaP cell lysate using an IgG or PTGES3 antibody. KAT2A, P300, FOXA1, and PTGES3 protein levels were detected by western blotting. **e,** LNCaP cells containing ARE-luciferase reporter were transfected with pCDH-PTGES3. Cells were treated with siRNA targeting control or KAT2A in the condition of 1nM DHT. Luciferase activities over Renilla were normalized with control (n = 3 as biological replicates; Mean ± SEM). Unpaired two-tailed t-test was used to determine statistical significance (**P < 0.01; ***P < 0.001).

**Extended data figure 8.**
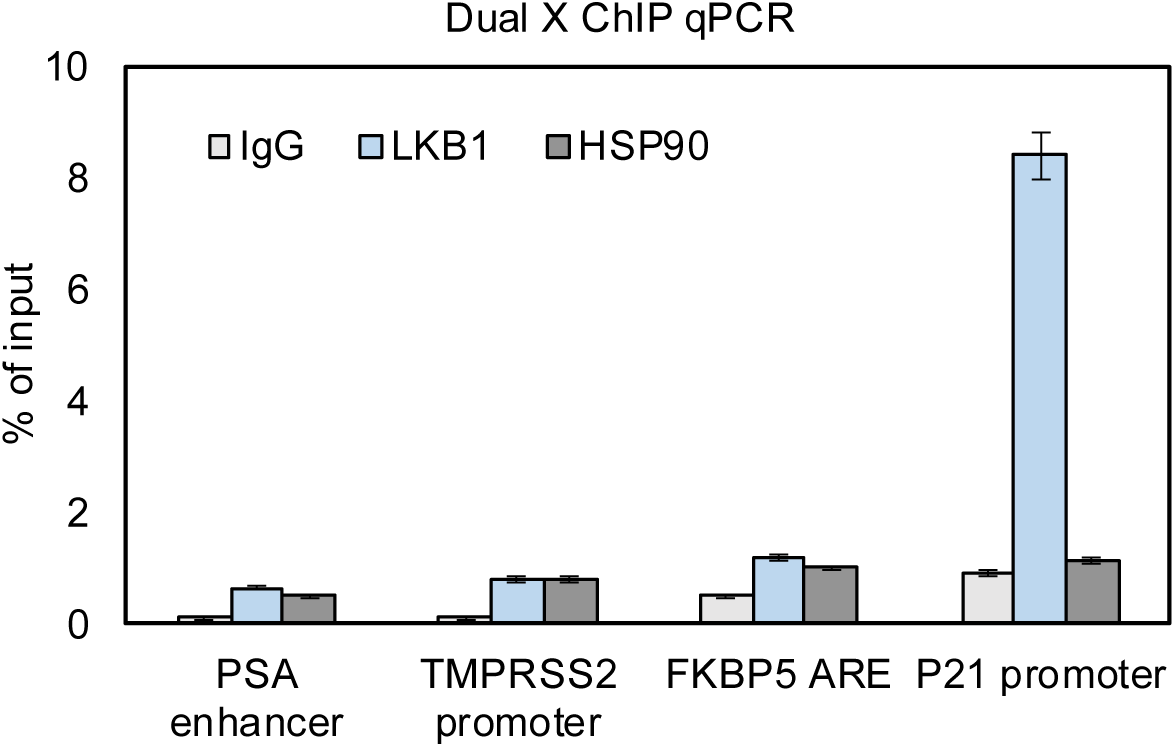
PTGES3 facilitates AR mediated transcription through a HSP90 independent manner. LNCaP cells were fixed sequentially by EGS and formaldehyde. Dual cross-linking ChIP experiments were performed using indicated antibodies. Precipitated DNA was used as a template to amplify the indicated genomic regions by real-time PCR (n = 3 as biological replicates; Mean ± SEM). LKB1 serves as a positive Dual X ChIP control which indirectly binds to the P21 promoter (non-ARE) by interacting with P53.

**Extended data figure 9.**
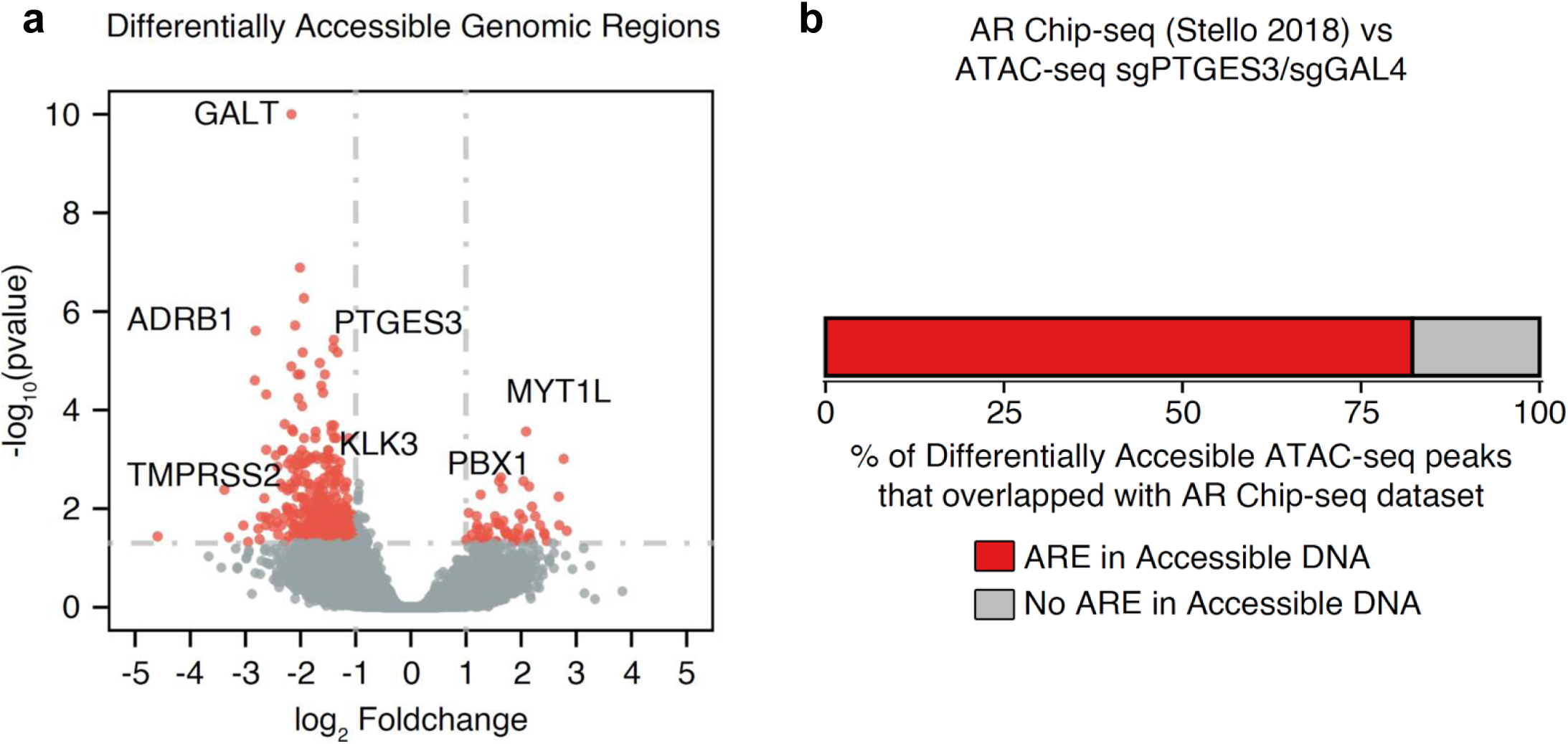
PTGES3 promotes the chromatin opening status of AR binding to AREs. **a,** LNCaPi cells were infected with sgGAL4 or sgPTGES3. Cells were collected for ATAC-Sequencing. The volcano plot shows the differentially accessible ATAC-seq peak regions in sgPTGES3 as compared to that in sgGAL4. Each dot represents an individual ATAC-seq peak. Differentially accessible ATAC-seq peaks are highlighted in red color (pvalue <0.05 and log2 Foldchange > 1 or <-1). Nearest gene to several representative differentially accessible ATAC-seq peaks have been highlighted. **b,** These differentially accessible ATAC-seq peak regions were compared against AR Chip-seq data in LNCaP cells (Stello2018). The figure shows the percentage of these differentially accessible ATAC-seq peaks that overlapped with the AR Chip-seq dataset.

**Extended data figure 10.**
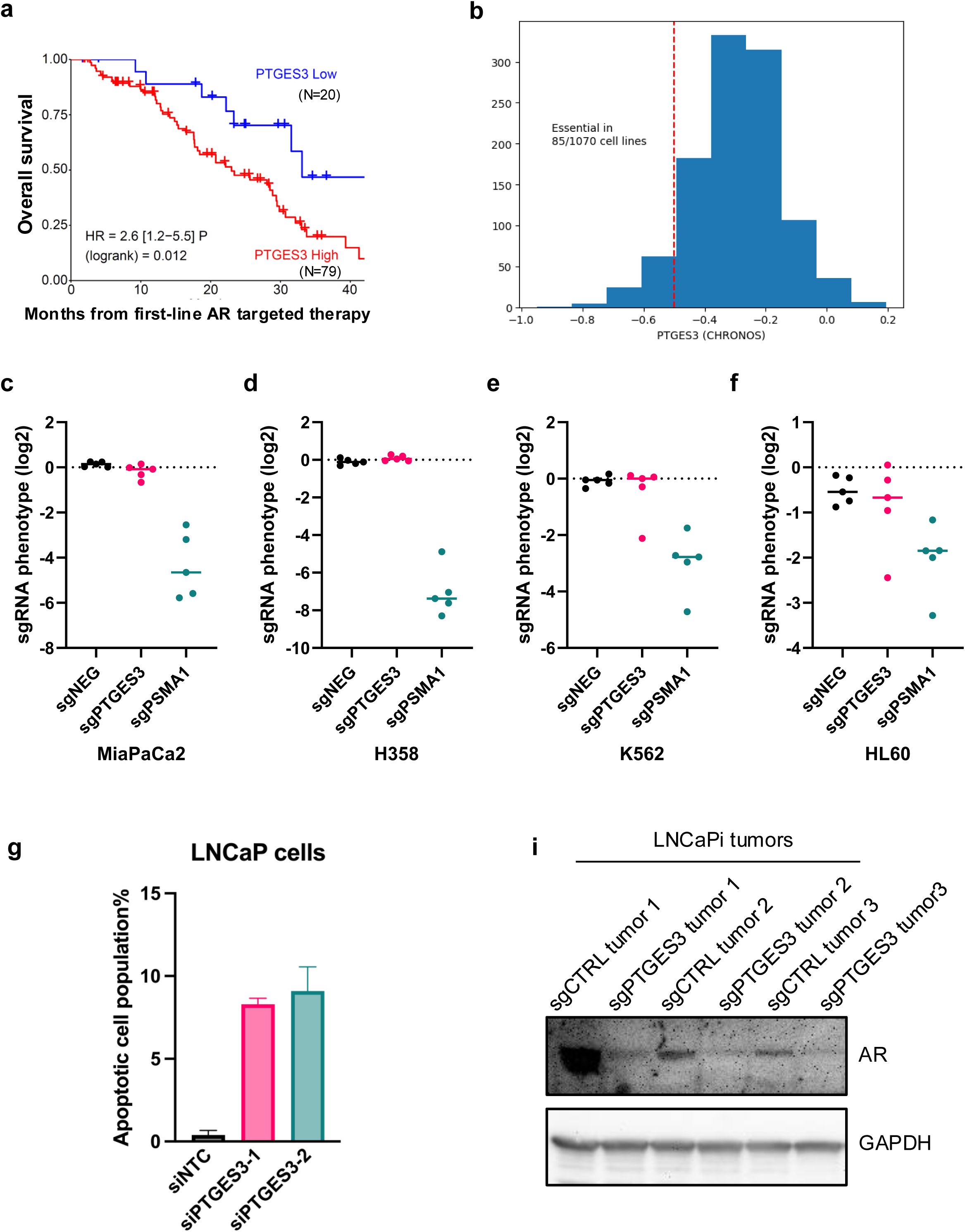
High PTGES3 expression predicts a poor outcome in advanced PCa patients treated with first line AR targeted therapy. **a**, Kaplan-Meier analysis of mCRPC patients treated with a first-line AR targeted therapy (abiraterone acetate, enzalutamide, or apalutamide) shows a worse overall survival in patients with high PTGES3 expression compared to low expression. **b**, A histogram of pan-cancer essentiality CHRONOS scores from the Cancer Dependency map showing the distribution of PTGES3 essentiality in 1070 cell lines. PTGES3 is considered essential for cell proliferation or survival in 85 cell lines (∼10% of all cell lines) with scores less than -0.5. This result indicates PTGES3 is not required for the vast majority of non-prostate cancer or AR independent cell lines. **c-f**, CRISPRi sgRNA level screen phenotypes performed in MiaPaCa2(**c**), H358(**d**), K562(**e**), and HL60(**f**) cells were plotted as log2 values for PTGES3, for PSMA1 which is a common essential gene and for 5 randomly selected non-targeting sgRNA controls present in the sgRNA library. **g**, LNCaP cells were treated with siRNA targeting control or PTGES3 for 48 hours, then stained with PI/Annexin V. The percentage of the apoptotic population was measured by FACS (n = 3 biological replicates; Mean ± SEM). **i**, Tumor tissues from Fig.4i were collected for western blot using AR and GAPDH antibodies.

## References

1. Robinson, D. et al. Integrative clinical genomics of advanced prostate cancer. Cell 161, 1215–1228 (2015).

2. Quigley, D. A. et al. Genomic Hallmarks and Structural Variation in Metastatic Prostate Cancer. Cell 174, 758–769.e9 (2018).

3. Attard, G. et al. Characterization of ERG, AR and PTEN gene status in circulating tumor cells from patients with castration-resistant prostate cancer. Cancer Res. 69, 2912–2918 (2009).

4. Ruizeveld de Winter, J. A., et al. Androgen receptor status in localized and locally progressive hormone refractory human prostate cancer. Am. J. Pathol. 144, 735–746 (1994).

5. Sadi, M. V., Walsh, P. C. & Barrack, E. R. Immunohistochemical study of androgen receptors in metastatic prostate cancer. Comparison of receptor content and response to hormonal therapy. Cancer 67, 3057–3064 (1991).

6. Culig, Z. & Santer, F. R. Androgen receptor signaling in prostate cancer. Cancer Metastasis Rev. 33, 413–427 (2014).

7. Pomerantz, M. M. et al. The androgen receptor cistrome is extensively reprogrammed in human prostate tumorigenesis. Nat. Genet. 47, 1346–1351 (2015).

8. Parolia, A. et al. Distinct structural classes of activating FOXA1 alterations in advanced prostate cancer. Nature 571, 413–418 (2019).

9. Smith, M. R. et al. Apalutamide Treatment and Metastasis-free Survival in Prostate Cancer. N. Engl. J. Med. 378, 1408–1418 (2018).

10. Beer, T. M. et al. Enzalutamide in metastatic prostate cancer before chemotherapy. N. Engl. J. Med. 371, 424–433 (2014).

11. Ryan, C. J. et al. Abiraterone in metastatic prostate cancer without previous chemotherapy. N. Engl. J. Med. 368, 138–148 (2013).

12. Bolla, M. et al. Long-term results with immediate androgen suppression and external irradiation in patients with locally advanced prostate cancer (an EORTC study): a phase III randomised trial. Lancet Lond. Engl. 360, 103–106 (2002).

13. Shipley, W. U. et al. Radiation with or without Antiandrogen Therapy in Recurrent Prostate Cancer. N. Engl. J. Med. 376, 417–428 (2017).

14. Chi, K. N. et al. Apalutamide for Metastatic, Castration-Sensitive Prostate Cancer. N. Engl. J. Med. 381, 13–24 (2019).

15. Davis, I. D. et al. Enzalutamide with Standard First-Line Therapy in Metastatic Prostate Cancer. N. Engl. J. Med. 381, 121–131 (2019).

16. James, N. D. et al. Abiraterone for Prostate Cancer Not Previously Treated with Hormone Therapy. N. Engl. J. Med. 377, 338–351 (2017).

17. Sternberg, C. N. et al. Enzalutamide and Survival in Nonmetastatic, Castration-Resistant Prostate Cancer. N. Engl. J. Med. 382, 2197–2206 (2020).

18. Fizazi, K. et al. Darolutamide in Nonmetastatic, Castration-Resistant Prostate Cancer. N. Engl. J. Med. 380, 1235–1246 (2019).

19. Watson, P. A., Arora, V. K. & Sawyers, C. L. Emerging mechanisms of resistance to androgen receptor inhibitors in prostate cancer. Nat. Rev. Cancer 15, 701–711 (2015).

20. Aggarwal, R. et al. Clinical and Genomic Characterization of Treatment-Emergent Small-Cell Neuroendocrine Prostate Cancer: A Multi-institutional Prospective Study. J. Clin. Oncol. Off. J. Am. Soc. Clin. Oncol. 36, 2492–2503 (2018).

21. Bluemn, E. G. et al. Androgen Receptor Pathway-Independent Prostate Cancer Is Sustained through FGF Signaling. Cancer Cell 32, 474–489.e6 (2017).

22. Visakorpi, T. et al. In vivo amplification of the androgen receptor gene and progression of human prostate cancer. Nat. Genet. 9, 401–406 (1995).

23. Grasso, C. S. et al. The mutational landscape of lethal castration-resistant prostate cancer. Nature 487, 239–243 (2012).

24. Takeda, D. Y. et al. A Somatically Acquired Enhancer of the Androgen Receptor Is a Noncoding Driver in Advanced Prostate Cancer. Cell 174, 422–432.e13 (2018).

25. Viswanathan, S. R. et al. Structural Alterations Driving Castration-Resistant Prostate Cancer Revealed by Linked-Read Genome Sequencing. Cell 174, 433–447.e19 (2018).

26. Gregory, C. W., Johnson, R. T., Mohler, J. L., French, F. S. & Wilson, E. M. Androgen receptor stabilization in recurrent prostate cancer is associated with hypersensitivity to low androgen. Cancer Res. 61, 2892–2898 (2001).

27. Henzler, C. et al. Truncation and constitutive activation of the androgen receptor by diverse genomic rearrangements in prostate cancer. Nat. Commun. 7, 13668 (2016).

28. Dehm, S. M., Schmidt, L. J., Heemers, H. V., Vessella, R. L. & Tindall, D. J. Splicing of a novel androgen receptor exon generates a constitutively active androgen receptor that mediates prostate cancer therapy resistance. Cancer Res. 68, 5469–5477 (2008).

29. Chang, K.-H. et al. A gain-of-function mutation in DHT synthesis in castration-resistant prostate cancer. Cell 154, 1074–1084 (2013).

30. Cabantous, S., Terwilliger, T. C. & Waldo, G. S. Protein tagging and detection with engineered self-assembling fragments of green fluorescent protein. Nat. Biotechnol. 23, 102–107 (2005).

31. Waldo, G. S., Standish, B. M., Berendzen, J. & Terwilliger, T. C. Rapid protein-folding assay using green fluorescent protein. Nat. Biotechnol. 17, 691–695 (1999).

32. Kamiyama, D. et al. Versatile protein tagging in cells with split fluorescent protein. Nat. Commun. 7, 11046 (2016).

33. Feng, S. et al. Improved split fluorescent proteins for endogenous protein labeling. Nat. Commun. 8, 370 (2017).

34. Leonetti, M. D., Sekine, S., Kamiyama, D., Weissman, J. S. & Huang, B. A scalable strategy for high-throughput GFP tagging of endogenous human proteins. Proc. Natl. Acad. Sci. U. S. A. 113, E3501–3508 (2016).

35. Horlbeck, M. A. et al. Compact and highly active next-generation libraries for CRISPR-mediated gene repression and activation. eLife 5, e19760 (2016).

36. Stelloo, S. et al. Endogenous androgen receptor proteomic profiling reveals genomic subcomplex involved in prostate tumorigenesis. Oncogene 37, 313–322 (2018).

37. Szklarczyk, D. et al. STRING v11: protein-protein association networks with increased coverage, supporting functional discovery in genome-wide experimental datasets. Nucleic Acids Res. 47, D607–D613 (2019).

38. De Leon, J. T. et al. Targeting the regulation of androgen receptor signaling by the heat shock protein 90 cochaperone FKBP52 in prostate cancer cells. Proc. Natl. Acad. Sci. 108, 11878–11883 (2011).

39. Eftekharzadeh, B. et al. Hsp70 and Hsp40 inhibit an inter-domain interaction necessary for transcriptional activity in the androgen receptor. Nat. Commun. 10, 3562 (2019).

40. Heinlein, C. A. & Chang, C. Androgen Receptor in Prostate Cancer. Endocr. Rev. 25, 276–308 (2004).

41. Reebye, V. et al. Role of the HSP90-associated cochaperone p23 in enhancing activity of the androgen receptor and significance for prostate cancer. Mol. Endocrinol. Baltim. Md 26, 1694–1706 (2012).

42. Cano, L. Q. et al. The co-chaperone p23 promotes prostate cancer motility and metastasis. Mol. Oncol. 9, 295–308 (2015).

43. Freeman, B. C. & Yamamoto, K. R. Disassembly of transcriptional regulatory complexes by molecular chaperones. Science 296, 2232–2235 (2002).

44. Knoblauch, R. & Garabedian, M. J. Role for Hsp90-associated cochaperone p23 in estrogen receptor signal transduction. Mol. Cell. Biol. 19, 3748–3759 (1999).

45. Tanioka, T., Nakatani, Y., Semmyo, N., Murakami, M. & Kudo, I. Molecular Identification of Cytosolic Prostaglandin E2 Synthase That Is Functionally Coupled with Cyclooxygenase-1 in Immediate Prostaglandin E2Biosynthesis. J. Biol. Chem. 275, 32775–32782 (2000).

46. Penning, T. M. AKR1C3 (type 5 17β-hydroxysteroid dehydrogenase/prostaglandin F synthase): Roles in malignancy and endocrine disorders. Mol. Cell. Endocrinol. 489, 82–91 (2019).

47. Jeong, K. H., Jung, J. H., Kim, J. E. & Kang, H. Prostaglandin D2-Mediated DP2 and AKT Signal Regulate the Activation of Androgen Receptors in Human Dermal Papilla Cells. Int. J. Mol. Sci. 19, (2018).

48. Zelin, E., Zhang, Y., Toogun, O. A., Zhong, S. & Freeman, B. C. The p23 molecular chaperone and GCN5 acetylase jointly modulate protein-DNA dynamics and open chromatin status. Mol. Cell 48, 459–470 (2012).

49. Stelloo, S. et al. Integrative epigenetic taxonomy of primary prostate cancer. Nat. Commun. 9, 4900 (2018).

50. Tsherniak, A. et al. Defining a Cancer Dependency Map. Cell 170, 564–576.e16 (2017).

51. Ge, A. Y. et al. A multiomics approach reveals RNA dynamics promote cellular sensitivity to DNA hypomethylation. Preprint at 10.1101/2022.12.14.518457 (2022).

52. Lou, K. et al. IFITM proteins assist cellular uptake of diverse linked chemotypes. Science 378, 1097– 1104 (2022).

53. Lou, K. et al. KRASG12C inhibition produces a driver-limited state revealing collateral dependencies. Sci. Signal. 12, eaaw9450 (2019).

54. Li, W. et al. MAGeCK enables robust identification of essential genes from genome-scale CRISPR/Cas9 knockout screens. Genome Biol. 15, 554 (2014).

55. Kothari, V. et al. DNA-Dependent Protein Kinase Drives Prostate Cancer Progression through Transcriptional Regulation of the Wnt Signaling Pathway. Clin. Cancer Res. Off. J. Am. Assoc. Cancer Res. 25, 5608–5622 (2019).

56. Roggero, C. M. et al. A detailed characterization of stepwise activation of the androgen receptor variant 7 in prostate cancer cells. Oncogene 40, 1106–1117 (2021).

57. Kong, A. T., Leprevost, F. V., Avtonomov, D. M., Mellacheruvu, D. & Nesvizhskii, A. I. MSFragger: ultrafast and comprehensive peptide identification in mass spectrometry-based proteomics. Nat. Methods 14, 513–520 (2017).

58. Barpanda, A. et al. Integrative Proteomic and Pharmacological Analysis of Colon Cancer Reveals the Classical Lipogenic Pathway with Prognostic and Therapeutic Opportunities. J. Proteome Res. 22, 871–884 (2023).

59. Mehra, R. et al. A novel RNA in situ hybridization assay for the long noncoding RNA SChLAP1 predicts poor clinical outcome after radical prostatectomy in clinically localized prostate cancer. Neoplasia N. Y. N 16, 1121–1127 (2014).

60. Schiewer, M. J. et al. PARP-1 regulates DNA repair factor availability. EMBO Mol. Med. 10, (2018).

61. Zeng, P.-Y., Vakoc, C. R., Chen, Z.-C., Blobel, G. A. & Berger, S. L. In vivo dual cross-linking for identification of indirect DNA-associated proteins by chromatin immunoprecipitation. BioTechniques 41, 694, 696, 698 (2006).

62. Kechin, A., Boyarskikh, U., Kel, A. & Filipenko, M. cutPrimers: A New Tool for Accurate Cutting of Primers from Reads of Targeted Next Generation Sequencing. J. Comput. Biol. J. Comput. Mol. Cell Biol. 24, 1138–1143 (2017).

63. Dobin, A. et al. STAR: ultrafast universal RNA-seq aligner. Bioinforma. Oxf. Engl. 29, 15–21 (2013).

64. Liao, Y., Smyth, G. K. & Shi, W. The R package Rsubread is easier, faster, cheaper and better for alignment and quantification of RNA sequencing reads. Nucleic Acids Res. 47, e47 (2019).

65. Love, M. I., Huber, W. & Anders, S. Moderated estimation of fold change and dispersion for RNA-seq data with DESeq2. Genome Biol. 15, 550 (2014).

66. Buenrostro, J. D., Giresi, P. G., Zaba, L. C., Chang, H. Y. & Greenleaf, W. J. Transposition of native chromatin for fast and sensitive epigenomic profiling of open chromatin, DNA-binding proteins and nucleosome position. Nat. Methods 10, 1213–1218 (2013).

67. Corces, M. R. et al. An improved ATAC-seq protocol reduces background and enables interrogation of frozen tissues. Nat. Methods 14, 959–962 (2017).

68. Martin, M. Cutadapt removes adapter sequences from high-throughput sequencing reads. EMBnet.journal 17, 10–12 (2011).

69. Andrews, S. FastQC: A Quality Control Tool for High Throughput Sequence Data. http://www.bioinformatics.babraham.ac.uk/projects/fastqc/ (2010).

70. Langmead, B. & Salzberg, S. L. Fast gapped-read alignment with Bowtie 2. Nat. Methods 9, 357–9 (2012).

71. Li, H. et al. The Sequence Alignment/Map format and SAMtools. Bioinformatics 25, 2078–2079 (2009).

72. Ramírez, F. et al. deepTools2: a next generation web server for deep-sequencing data analysis. Nucleic Acids Res. 44, W160–5 (2016).

73. Zhang, Y. et al. Model-based analysis of ChIP-Seq (MACS). Genome Biol. 9, R137 (2008).

74. Yu, G., Wang, L.-G. & He, Q.-Y. ChIPseeker: an R/Bioconductor package for ChIP peak annotation, comparison and visualization. Bioinforma. Oxf. Engl. 31, 2382–3 (2015).

75. Bailey, T. L. et al. MEME SUITE: tools for motif discovery and searching. Nucleic Acids Res. 37, W202–8 (2009).

76. Fornes, O. et al. JASPAR 2020: update of the open-access database of transcription factor binding profiles. Nucleic Acids Res. 48, D87–D92 (2020).

77. Karnes, R. J. et al. Development and Validation of a Prostate Cancer Genomic Signature that Predicts Early ADT Treatment Response Following Radical Prostatectomy. Clin. Cancer Res. Off. J. Am. Assoc. Cancer Res. 24, 3908–3916 (2018).

78. Abida, W. et al. Genomic correlates of clinical outcome in advanced prostate cancer. Proc. Natl. Acad. Sci. 116, 11428–11436 (2019).

79. Cerami, E. et al. The cBio Cancer Genomics Portal: An Open Platform for Exploring Multidimensional Cancer Genomics Data: Figure 1. Cancer Discov. 2, 401–404 (2012).

80. Gao, J. et al. Integrative Analysis of Complex Cancer Genomics and Clinical Profiles Using the cBioPortal. Sci. Signal. 6, pl1–pl1 (2013).

81. Therneau, T. M. A Package for Survival Analysis in R. (2021).

82. Kassambara, A., Kosinski, M. & Biecek, P. Survminer: Drawing Survival Curves Using ‘Ggplot2’. R Package Version 0.*4*.*9*. (2021).

83. Das, R. et al. An integrated functional and clinical genomics approach reveals genes driving aggressive metastatic prostate cancer. Nat. Commun. 12, 4601 (2021).

84. Hieronymus, H. et al. Gene expression signature-based chemical genomic prediction identifies a novel class of HSP90 pathway modulators. Cancer Cell 10, 321–330 (2006).

